# Phages against non-capsulated *Klebsiella pneumoniae*: broader host range, slower resistance

**DOI:** 10.1101/2022.08.04.502604

**Authors:** Marta Lourenço, Lisa Osbelt, Virginie Passet, François Gravey, Till Strowig, Carla Rodrigues, Sylvain Brisse

## Abstract

**Background:** *Klebsiella pneumoniae* (Kp) is an ecologically generalist bacterium but also an opportunistic pathogen responsible for hospital-acquired infections and a major contributor to the global burden of antimicrobial resistance. In the last decades, few advances have been made in the use of virulent phages as alternative or complement to antibiotics to treat Kp infections. The efficiency of phages relies on their ability to recognize and attach to the bacterial surface structure, and in the case of Kp, capsule (K) is the main surface structure. However, Kp capsule is highly polymorphic and the majority of classically isolated phages are specific for unique K-types, limiting therapy prospects. In this study, we demonstrate the feasibility of an innovative strategy consisting in isolating phages that target capsule-deficient mutant Kp strains, and compare such phages with anti-capsulated cells phages phylogenetically and through *in vitro* and *in vivo* experiments.

**Methods:** We isolated 27 phages using 7 capsule-deficient Kp strains as hosts (anti-K^d^ phages), and 41 phages against 7 wild-type (wt) Kp strains (anti-K phages). We evaluated and compared phenotypically and genotypically their host range, resistance emergence and selected mutations and *in-vivo* activity.

**Results:** *In vitro*, anti-K^d^ phages showed a broader host-range, with most phages being able to infect non-capsulated mutants of multiple sublineages and O-antigen locus types. Besides, the emergence of bacterial subpopulations non-susceptible to anti-K^d^ phages was slower when compared to anti-K phages and with a different range of genomic differences. One anti-K^d^ phage (mtp5) was shown to infect non-capsulated Kp strains belonging to 10 of the 12 known O-antigen types. Moreover, this phage was able to replicate in the gut of mice colonised with the wt (capsulated) parent strain.

**Conclusions:** This work demonstrates the potential value of an anti-*Klebsiella* phage isolation strategy that addresses the issue of narrow host-range of anti-K phages. Anti K^d^-phages may be active in infection sites where capsule expression is intermittent or repressed, or in combination with anti-K phages, which often induce loss of capsule escape mutants.

## Introduction

*Klebsiella pneumoniae* (Kp), a bacterium found in a wide variety of ecological compartments (soil, plants, water), is a common gut colonizer of humans and animals, but also an important opportunistic pathogen. Kp is responsible for a broad range of infections, including urinary tract or respiratory infections, liver abscess and septicaemia and is a major contributor to the global burden of antimicrobial resistance (AMR) (WyresNatR2020, Antimicrobial Resistance Collaborators 2022). The current increase of multidrug resistant (MDR) Kp infections, coupled with the growing number of reports of pandrug-resistant Kp strains (Rodrigues et al. 2022) untreatable by current antimicrobial therapy, as well as the increase in the number of invasive infections by hypervirulent Kp, has renewed the interest in Kp biology with the prospect of developing alternative therapies that may be complementary to antimicrobials.

One of the approaches that has raised more interest over the years relies on the use of virulent bacteriophages (known as phage therapy), which are viruses that target and kill bacteria (Roach and Debarbieux 2017). The killing efficacy of phages relies, among others, on their ability to attach to the bacterial surface, implying a high specificity towards surface structures. In Kp, the most external structure is the capsule (K antigen), a thick layer of polysaccharides that surrounds the bacteria and masks other cell wall structures, such as the lipopolysaccharide (LPS) and its O-antigen. This makes the capsule the primary receptor for most phages that have been isolated against Kp up to now (C. R. Hsu et al. 2011; Hoyles et al. 2015; Gorodnichev et al. 2021; Hao et al. 2021; Bonilla et al. 2021; Eckstein et al. 2021; Pertics et al. 2021; Townsend et al. 2021; Beamud et al. 2022). With the exception of hypervirulent Kp infections, which are generally associated with a narrow range of KL-types (mainly KL1 and KL2) (Lin et al. 2014; Hung et al. 2011), capsular structure polymorphism is high in Kp, and varies even within narrow phylogenetic clonal groups (Rodrigues et al. 2020; Wyres et al. 2019; Rodrigues et al. 2022; Lam, Wick, Watts, et al. 2021). Using serological methods, 77 distinct K-types (K1 to K82) have been distinguished (Ørskov and Ørskov 1984), and genomic sequencing has uncovered many additional capsular polysaccharide synthesis (*cps*) loci (KL-types, KL101-KL186) (Lam, Wick, Watts, et al. 2021; Lam, Wick, Judd, et al. 2021), which likely synthesize yet distinct capsular structures. The distribution of KL types among clinical isolates of Kp shows that the six most frequent K-types (KL107, KL2, KL24, KL106, KL64, KL17) only represent 38% of Kp isolates, whereas 17 additional K-types would be needed to reach 75% (Lam, Wick, Watts, et al. 2021).

In addition to the challenge of the high polymorphic Kp capsule, resistance to phages that attach to the capsule is also known to evolve relatively fast, and predominantly via loss of capsule production (Eckstein et al. 2021; Hesse et al. 2020; Fang et al. 2022; Majkowska-Skrobek et al. 2021). This phage escape mechanism thus results in the exposure of other membrane structures such as the LPS, O-antigen or outer membrane proteins. Interestingly, O-types (8 types serologically defined and 4 putative other ones defined by genomic analysis) and other surface Kp proteins are less diverse than capsular structures, representing potential broad-range phage targets. For example, types OL1, OL2 and OL3b represent 80% of Kp human infections (Lam, Wick, Watts, et al. 2021; Lam, Wick, Judd, et al. 2021). A rarely considered possibility is that *in vivo*, the capsular polysaccharide may not always be present. Recent studies have pointed out that while hypercapsulated phenotypes seem to be important for enhanced Kp dissemination through the bloodstream, capsule production is associated with a fitness cost in the gut (Y. H. Tan et al. 2020). Moreover, capsule loss seems to be beneficial for epithelial cell invasion and linked with persistent urinary tract infections (Ernst et al. 2020).

We hypothesized that phages attaching to molecular structures of the cell surface lying below the capsule, here collectively referred as anti-K^d^ (for anti-capsular deficient) phages, may represent interesting novel components of anti-*Klebsiella* strategies. Such phages have already been isolated (Hesse et al. 2020; Majkowska-Skrobek et al. 2021) but their broad-range potential and differences from anti-capsule (anti-K) phages have yet to be uncovered. The use of anti-K^d^ phages in combination with anti-K phages may also address the issue of phage resistance emergence when used as therapy, given that simultaneous modifications in different membrane structures would require additional mutations and probably result in high fitness costs (Koskella et al. 2012). This strategy may concomitantly cover the issue of the existence of multiple Kp populations *in vivo* (capsulated and non-capsulated strains).

Here, we isolated, characterized and compared *in silico, in vitro* and *in vivo*, anti-K^d^ phages with anti-K phages, and their interactions with their bacterial hosts. We show that anti-K^d^ phages have a broad host range and induce a slower rate of emergence of resistance *in vitro*, and are synergistic with anti-K phages. In particular, we discovered phage mtp5, which is able *in vitro* to lyse nearly all non-capsulated Kp strains tested. *In vivo*, this anti-K^d^ phage was able to replicate in the mice gut colonized with the wild-type (capsulated) Kp strain.

## Results

### Selection of wild-type and capsule-deficient strains for isolation of anti-K and anti-K^d^ phages

Multilocus sequence typing (MLST) of 7,388 *K. pneumoniae* species complex (henceforth, Kp) public genomes revealed a high genetic diversity, with more than 1,193 Sequence types (STs), with 7 STs being predominant (ST258, ST11, ST15, ST512, ST101, ST307 and ST147) and representing 55% of all genomes (**Figure S1, TableS1; Supplementary text 1**). To eliminate bias due to recent clonal expansions or outbreaks, only one strain per ST was retained to analyse the frequency of capsular and the O-antigen locus types (KL and OL, respectively). KL1, KL64, KL2, KL10, KL30, KL102, KL106 and KL107 were the most prevalent KL-types, while OL1, OL2 and OL3/O3a were the predominant OL-types (**Figure S1**), together representing 80% of Kp genomes.

For anti-K phage isolation (**Table** 1; **Supplementary text** 1), 7 Kp wild-type (wt, i.e., capsulated) strains (**Table 1, Table S2**) representative of the major KL-types were selected as hosts. To perform anti-K^d^ phages isolation, we selected 7 Kp capsule-deficient strains derived by mutagenesis in a previous study (de Sousa et al. 2020) and which represented the three most prevalent OL-types.

### Isolation and genomic characterization of 68 new phages

We isolated and sequenced 68 new phages: 41 anti-K phages and 27 anti-K^d^ phages (**Table S3**; Hereafter, the prefixes cp (for capsule polysaccharide phages) and mtp (for mutant capsule polysaccharide phages) will be used to name anti-K phages and anti-K^d^ phages, respectively. Regarding the lifestyle, all the newly isolated phages were categorized as virulent phages (BACPHLIP) (Hockenberry and Wilke 2021; Mavrich and Hatfull 2017). No genes encoding for putative virulence factors or antibiotic resistance were found in any of these phages.

A phylogenetic analysis of the isolated phages was performed together with 98 publicly available complete anti-*Klebsiella* phage genomes. Our phages were distributed into five phage families: *Autographiviridae, Drexlerviridae, Myoviridae, Ackermannviridae* and *Schitoviridae* (**Figure 1**). Whereas the majority anti-K phages were associated with the *Autographiviridae* family (63%, 26/41), 19 out of 27 (70%) anti-K^d^ phages belonged to *Drexlerviridae* (**Figure 1, Table S3**). Despite the high similarity detected between certain phages (e.g. cp33-38 [ANI 99.9%]; or mtp6 and cp48 [ANI 99.9%]), we decided to keep them as unique phages since small nucleotide differences can result in in host-range differences (De Sordi, Khanna, and Debarbieux 2017), as observed for mtp6 and cp48, able to infect different phenotypes of the same host (capsule-deficient and wild-type, respectively).

**Figure 1.**
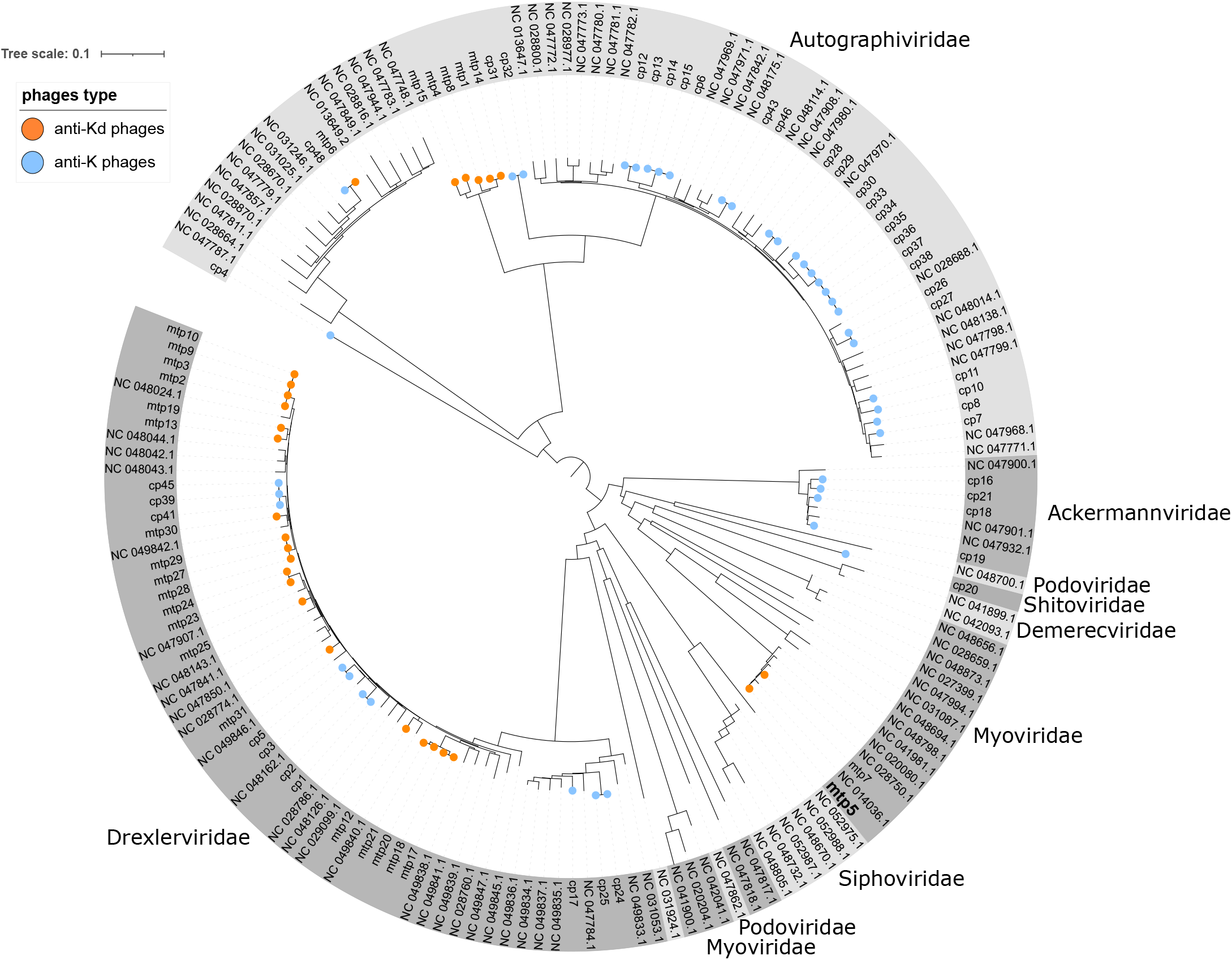
Phage diversity. Genome sequence-based phylogenetic tree of the phages isolated in this study, together with 98 other anti-*K*. *pneumoniae* phages (NCBI, March 2021). The phage families are indicated around the tree. Orange and blue circles at the tip of branches represent the phages isolated in this work: Orange: anti-K^d^ phages, Blue: anti-K phages. The tree is derived from a K-mer distance matrix (see Methods).

Several virulent anti-*Klebsiella* phages are known to have depolymerase functional domains in their tail fibers, giving them the ability to digest specific exopolysaccharides. Protein analysis and comparison with known depolymerases revealed between 0 and 5 depolymerase domains in the genomes of anti-K phages, whereas anti-K^d^ phages had only 0 or 1 domains (**Table S4**).

### Anti-K^d^ phages have a broader host range in comparison with anti-K phages

Based on a panel of 50 strains representing 31 KL-types, the anti-K phages targeted predominantly a unique KL-type (**Figure S2, Table S2**). Exceptions included (i) phages cp8-cp13 that infected KL10 and KL25 strains; (ii) phage cp34, which infected KL64 and KL102 strains; and (iii) phage cp16, which targeted KL35 and KL107 strains. Additionally, phages cp39, cp41 and cp45, which infected wt KL106 strains, also infected 4 out of 7 capsule-deficient mutant strains (**Figure S2, Table 1**) with the same OL-type (OL1v1). Interestingly, genomic sequencing revealed the presence of only one depolymerase domain for these three phages, which is homologous to a domain present in several anti-K^d^ phages.

In sharp contrast, anti-K^d^ phages infected a broader range of strains: each of them infected strains of at least 2 different OL-types (among the 3 OL-types of the 7 capsule-deficient mutant strains, which have 5 different KL-types) (**Figure 2, Figure S3**). As expected, anti-K^d^ phages were not active against the wt (capsulated) strains *in vitro*.

**Figure 2.**
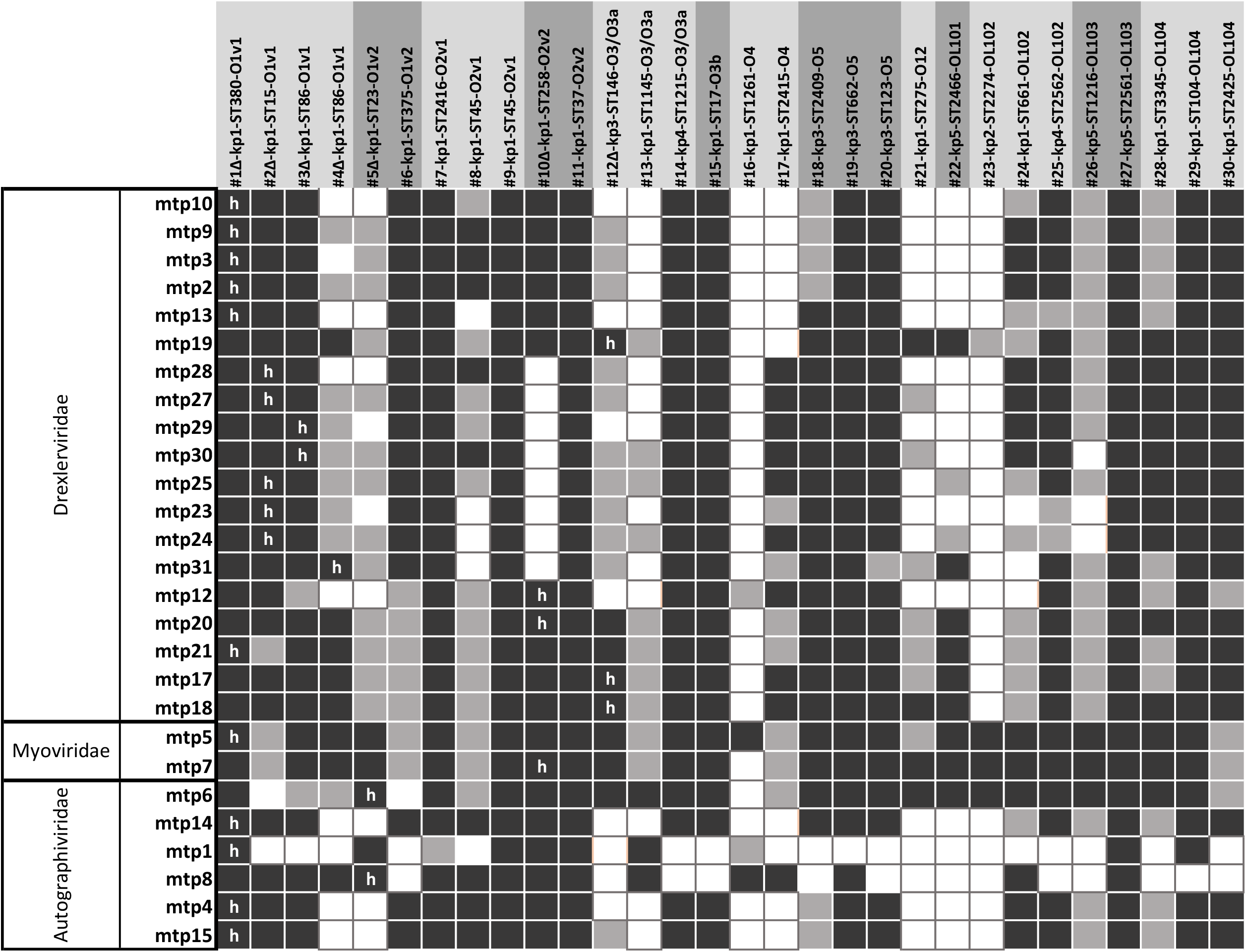
Host range of anti-K^d^ phages. The phages were tested against capsule-deficient strains (n=7, *wza* or *wcaJ* mutants, represented by a Δ) and spontaneous non-mucoid culture-derived strains (n=23). Dark orange: complete lysis; light orange: intermediate lysis; empty squares: absence of lysis. “h”: initial host used for phage isolation.

To investigate in more depth the host-range of anti-K^d^ phages, we generated 23 spontaneous non-mucoid clones from Kp strains belonging to 22 different STs, at least 18 KL-types and 11 different OL-types (out of the 12 currently known; the missing one is O8, of very low prevalence, representing only 0.03% of the 7,388 genomes analysed; **Table S2**) (Follador et al. 2016; Wick et al. 2018). Excepting mtp1 and mtp8, which only targeted 10 and 16 strains, respectively, the anti-K^d^ phages infected 20 to 28 strains out of the 30 non-capsulated (capsule-deficient or non-mucoid) strains (**Figure 2**). Among these, the broadest host-range phages were: (i) mtp5, which infected all the 30 strains; (ii) mtp7, mtp17, mtp18, mtp20 and mtp21 (28 strains each); and (iii) mtp6, mtp19 and mtp25 (27 strains each; **Figure 2**).

### Characteristics of mtp5, an anti-K^d^ phage with broad host-range

Phage mtp5 presented a 175.1 kb genome composed of 288 CDSs and 1 tRNA gene and belonged to the *Slopekvirus* genus within the *Myoviridae* family. Its genomic structure was closely related (ID ranging from 98.43% to 98.98%, and coverage of 94 to 97%) to phage mtp7 (this study) and other previously described anti-*Klebsiella* phages Matisse, Kp27, Kp15, Miro and PMBT1 (**Figure S4**) (Maciejewska et al. 2017; Provasek et al. 2015; Mijalis et al. 2015; Koberg et al. 2017). The main genomic difference between mtp5 and these related phages was found in one of the two tail fibers genes (**Figure S4**), the L-shaped tail fiber protein. The highest identity of this protein with the above-mentioned phages was with Miro phage (93.24%). When used as alone query against public sequence repositories, this protein showed 95.41% similarity with the L-shaped tail fiber protein of *Klebsiella* phage Kpn-VAC66 (GenBank accession no. MZ612130.1; a 178 Kb *Myoviridae* phage).

Efficiency of plating (EOP) assays confirmed the ability of mtp5 to infect all non-capsulated strains (**Figure 3A**). Further, when tested against wt strains in combination with anti-K phages (**Figure 3B**), phage mtp5 improved the lysis effect on all tested strains, with the single exception of strain 899 (ST101-KL106-OL1v2; **Figure 3B**).

**Figure 3.**
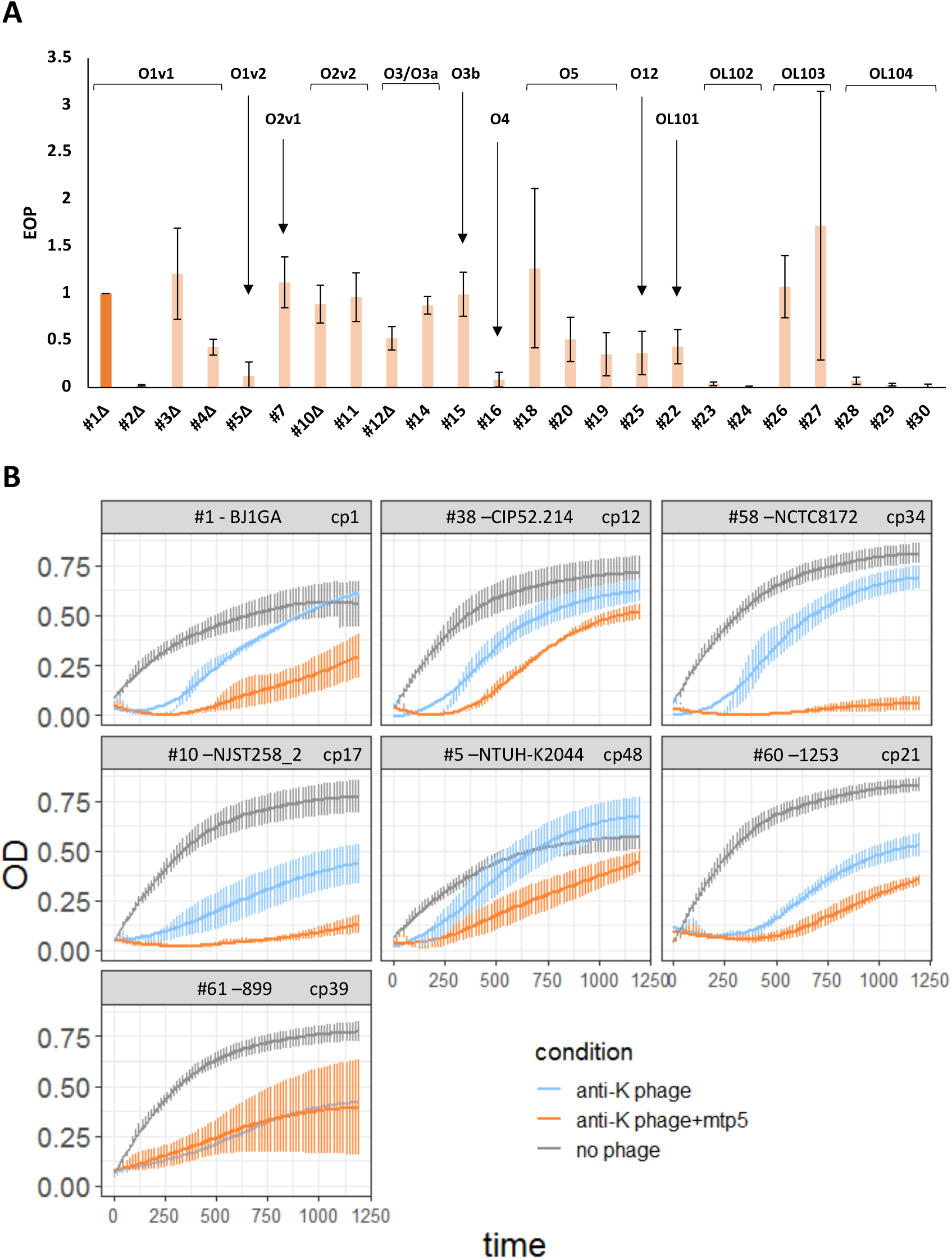
Efficacy of phage mtp5 on spontaneous non-mucoid mutants or in combination with anti-K phages. **A**. Efficiency of plating of phage mtp5 on non-mucoid spontaneous mutants. Bacterial strains are ordered by O-type as indicated and the EOP was calculated as a ratio of phage titers relative to the host strain, using 1 as the reference value of the host. In darker orange is represented the strain used for mtp5 phage isolation. Only 24 out of 30 non-capsulated strains are represented. For the remaining six strains, we were not able to calculate the infection titers, although we could still see lysis with the highest phage concentrations (These conditions might correspond to “lysis from without” in which high phage concentrations lead to bacterial lysis without actual infection and phage production (Abedon 2011)) **B**. Association of phage mtp5 with anti-K phages, assayed against capsulated strains. Growth curves are shown for the indicated Kp strains either without phage (grey) or with anti-K phages alone (blue) or in cocktail with mtp5 (orange; equal proportions of each). Three to five replicates were performed for each condition; the vertical error bars represent SEM (standard error of the mean). Phages were added at t=0 at MOI of 1 × 10E-2.

Other broad host-range anti-K^d^ phages were tested in combination with anti-K phages, and also showed increased killing effect and slow regrowth of escape mutants (see below), consistent with the observations made for mtp5 (**Figure S5**).

### Resistance against anti-K^d^ phages emerges slower than against anti-K phages

The emergence of subpopulations of bacteria that escape phage predation is typically observed in *in vitro* bacterial cultures (Hernandez and Koskella 2019; Hesse et al. 2020). We compared the timing of this phenomenon for anti-K phages and anti-K^d^ phages (**Figure S6**). With both categories of phages, the impact of phage predation became visible after 45 to 60 min post-infection. Interestingly, resurgence of growth was significantly delayed in bacterial cultures infected with anti-K^d^ phages when compared to those infected with anti-K phages (Area Under the Curve ratios: 0.2 to 0.4 vs 0.4 to 0.6, respectively; p-value = 0.006, Mann-Whitney test; **Figure S6C**). Simultaneously, we calculated the relative bacterial growth (RGB) at 4 h from cultures with anti-K or anti-K^d^ phages, and these were very low (from <0 to 0.2, p-value=0.4323) for the majority of strains, indicating the high efficiency of the both phages in controlling their target bacterial populations. However, after 6 h, 8 h and 10 h, the RGB of the non-susceptible populations was significantly lower for non-capsulated strains than for capsulated strains (*p*-value= 1.119×10^−06^, 2.204×10^−04^ and 3.332×10^−04^, respectively) (**Figure S6DE**). These results highlight the slower emergence of anti-K^d^ phages escape mutants, relative to anti-K phage escape ones.

### Genomic changes associated with anti-K and anti-K^d^ phages resistance

We next characterized and compared the bacterial genetic changes associated with anti-K and anti-K^d^ phage resistance. In the case of the non-susceptible population to anti-K phages, we detected mutations in 22 different genes, with the majority of them occurring in *cps* cluster genes, mainly in *wcaJ* or *wbaP* (**Figure 4, Figure S7, Table S5**). These mutations included single-nucleotide polymorphisms (SNPs), deletions and disruptions by insertion sequences (IS); and the relative frequency of these two events varied depending on the strains/phage combination. Interestingly, in two cases *wbaP* was disrupted by an IS91 family element, which was harboured by one of the plasmids of the strain (**Figure 4, Table S5**).

**Figure 4.**
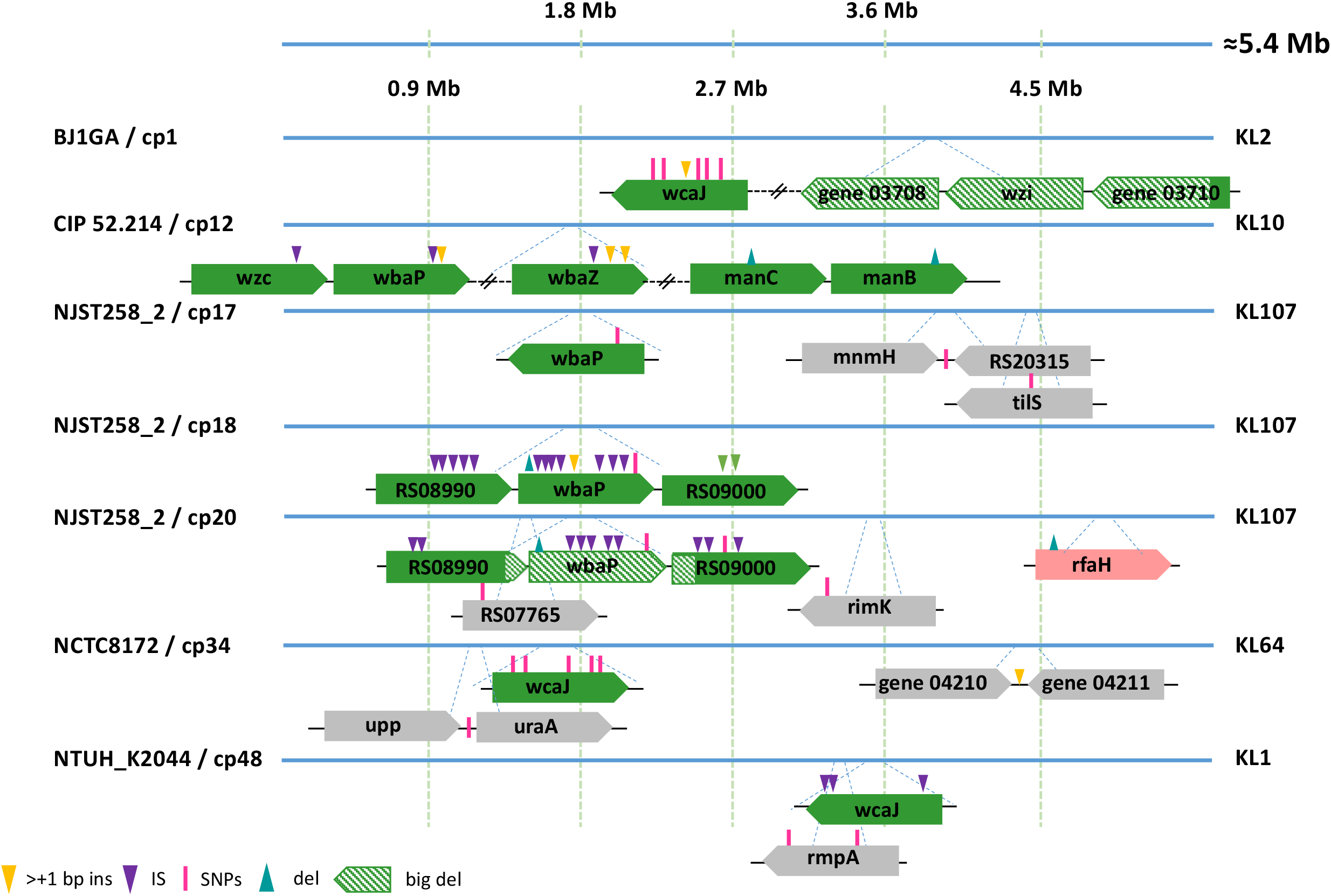
Mutations observed in populations exposed to anti-K phages. Clones were isolated after exposure to anti-K phages. Population sequence analysis (see main text) revealed the mutations depicted above. Genes representation colors: green: capsule locus, blue: O-antigen locus, pink: LPS locus and grey: others. For more details see table S7.

In contrast, resistance against anti-K^d^ phages was associated with more variable events (**Figure 5, Table S5**). Overall, mutations in 31 distinct genes from at least 12 different operons were found among anti-K^d^ resistant populations. Half (16/31) of these included mutations in (i) the O-antigen locus (*gspA, rfb* operon among others) and were observed in 4 out of 7 strain/phage combinations; (ii) the LPS biosynthesis genes, which were mutated in the combination of two anti-K^d^ phages with NJST258_2Δ*wza* strain; (iii) and surprisingly, in the *cps* locus (*wcaJ* and RS17345 genes), detected for the NTUH_K2044Δ*wza*/mtp6 combination (see additional information in **Supplementary text 2**). The remaining mutations mostly affected genes coding for transporters (ferrichrome^-^iron outer membrane transporter *fhuA, tonB* transport system; *uraA* coding for an uracil permease; *mntH* encoding a divalent metal cation transporter) or genes involved in carbohydrate metabolism (*purK* encoding a formate-dependent phosphoribosylglycinamide formyltransferase; *galE* coding for UDP-glucose 4-epimerase). Together, these results indicate that a variety of surface structures were disrupted in the phage resistant mutants. This diversity suggests that a wide set of receptors can be used by anti-K^d^ phages. We also noted that mutations in *rimK* (alpha-L-glutamate ligase), *uraA* and *upp* (uracil phosphoribosyltransferase) genes were associated both with anti-K and anti-K^d^ phage non-susceptible populations, suggesting they affect cellular processes involved in capsular and non-capsular receptors.

**Figure 5.**
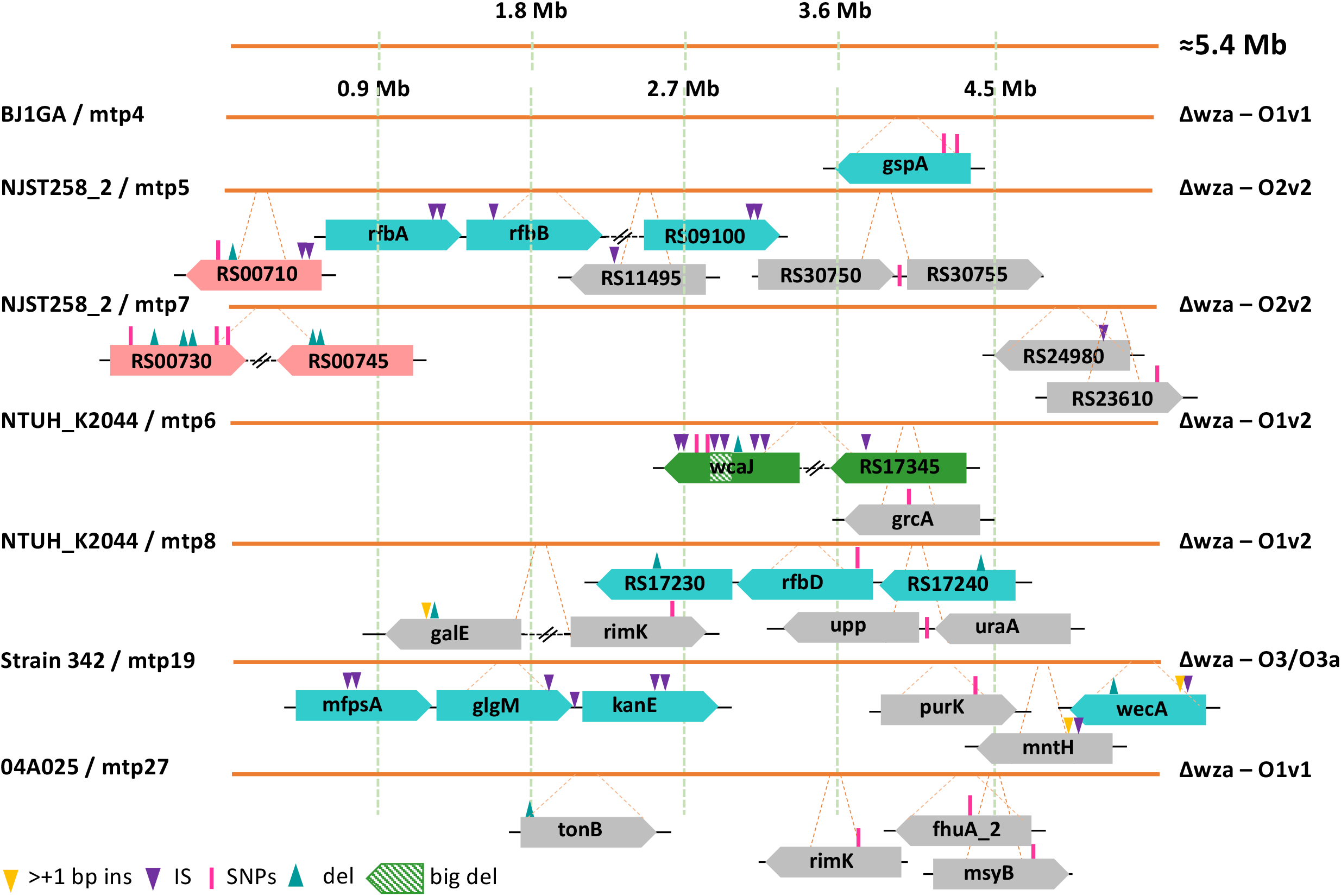
Mutations observed in populations resisting against anti-K^d^ phages. Clones were isolated after exposure to anti-K^d^ phages. Population sequence analysis (see main text) revealed the mutations depicted above. Genes representation colors: green: capsule locus, blue: O-antigen locus, pink: LPS locus and grey: others. For more details see table S6.

### Mutations associated with resistance against the broad-range anti-K^d^ phage mtp5

When mtp5 was used alone, mtp5-resistant populations of BJ1GAΔ*wza* (its original host) emerged after around 6.5 h, but no mutations could be detected (as assessed at 18 h of infection). We therefore used the NJST258_2Δ*wza*/mtp5 combination, and found that the non-susceptible population had mutations in several genes related to membrane biosynthesis, including in gene RS00710 encoding a glycosyltransferase involved in core LPS biosynthesis (**Figure 5, Table S5**).

We next tested four phage cocktails of mtp5 with anti-K phages (cp1, cp12, cp34 and cp48) against capsulated strains, and characterized the genetic mutations involved in the emergence of non-susceptible populations (**Figure 6**). In total, mutations in 8 different genes were detected. Among these were the *cps* initiator glycosyltransferase genes *wcaJ* (BJ1GA/cp1+mtp5 and NTUH_K2044/cp48+mtp5) and *wbaP* (CIP52.214/cp12+mtp5), which were also associated with resistance against the anti-K phages alone (**Figure 4**). The remaining mutations concerned mainly genes involved in carbohydrate (*galU, xylAB*) and lipids (*acpP*) metabolism. In the case of *galU* (encoding for α-D-glucose-1-phosphate uridylytransferase) a single mutation was detected in BJ1GA/cp1+mtp5 (this mutation also detected in BJ1GA/cp1+mtp4). Gene *galU* plays a central role in cell envelope synthesis and was previously associated with resistance to two different anti-*Klebsiella* phages, with impaired growth rate (Hesse et al. 2020). Regarding gene *acpP*, coding for an acyl carrier protein involved in fatty acid biosynthesis and in membrane-derived-oligosaccharide biosynthesis in *E. coli* (Rawlings and Cronan 1992; Keseler et al. 2011), a non-synonymous point mutation was found in NCTC8172/cp34+mtp5. For this bacteria/phage combination no genes from the *cps* cluster were affected, suggesting that mtp5 resistance mutations can be sufficient to prevent infection by the two phages (**Figure 6**). Finally, we next tested phage mtp5 in cocktail with a second anti-K^d^ phage, mtp4, against the capsule-deficient mutant strain BJ1GAΔ*wza*. The resistant population showed an 89 kb deletion, comprising 72 different genes encoding for capsule and O-antigen synthesis, and for some genes of the core LPS biosynthesis. Additionally, this large deletion also removed the histidine operon and disrupted genes *pksJ* (of the colibactin operon) and *dnaK*. These observations suggest that escape to combination of anti-K^d^ phages may incur large fitness costs, as they affect different pathways that are key to the bacterial physiology.

**Figure 6.**
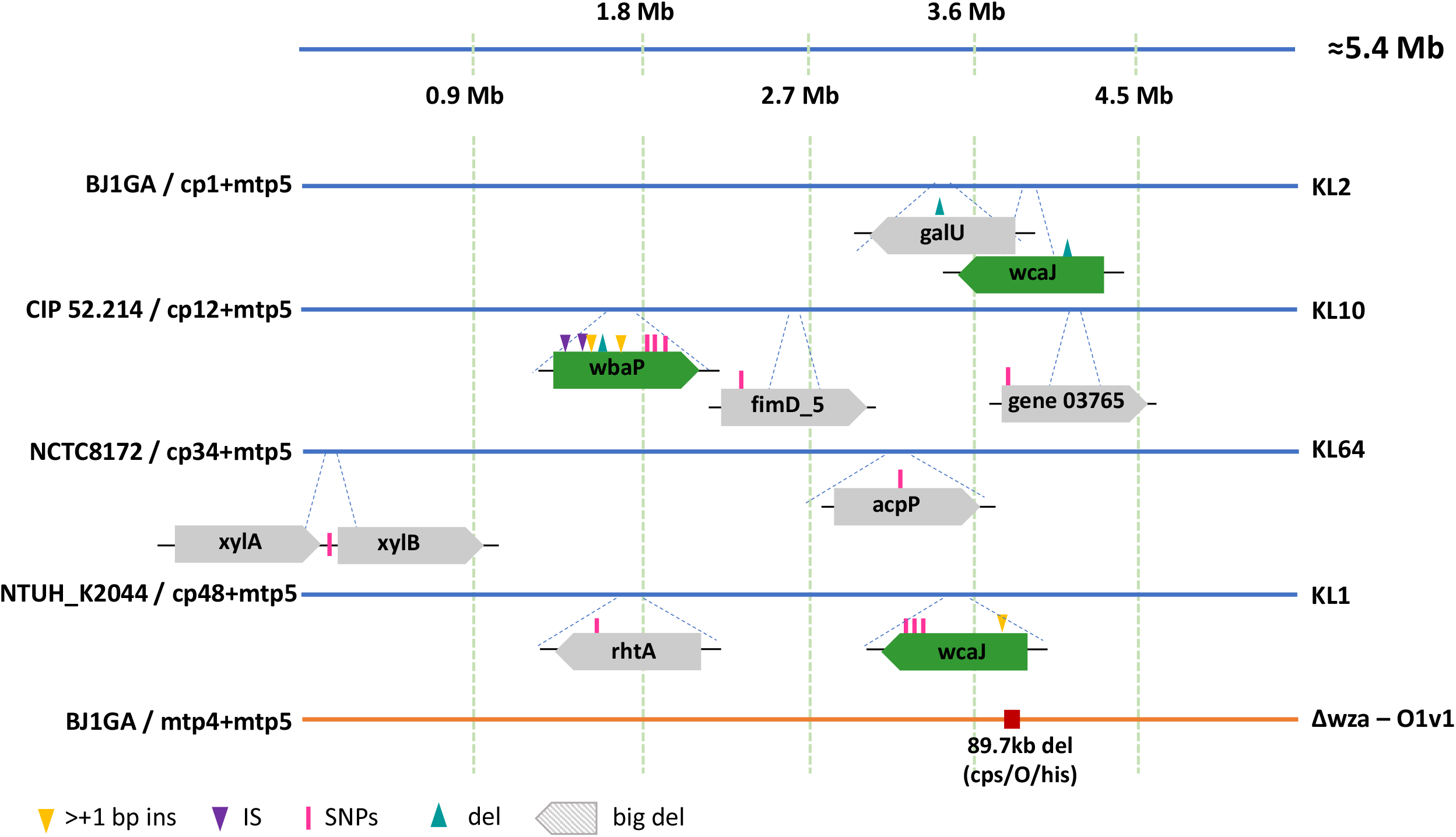
Mutations observed in populations resisting against anti-K+mtp5 phage cocktails. Clones were isolated after exposure to cocktails combining anti-K phages and the broad-range anti-K^d^ phage mtp5. Sequence analysis revealed the mutations depicted above (for more information see **Table S6**). Top (genomes represented by blue lines): wt strains; bottom (genomes represented by orange lines): capsule-deficient strains. Genes representation colors: green: capsule locus, blue: O-antigen locus, pink: LPS locus and grey: others.

Taken together, these results show that resistance to mtp5 is mainly linked to disruptions of the production of the initial building blocks of the LPS biosynthesis.

### Resistance to other phage cocktails

Resistance mutations were assessed for three other anti-K/anti-K^d^ phage cocktails (BJ1GA/cp1+mtp4, NJST258_2/cp17+mtp17 and NTUH_K2044/cp48+mtp6) (see **supplementary text 3, Figure S8**). In general, mutations observed in the non-susceptible populations concerned transporters and genes involved in LPS O-locus biosynthesis, similar to what was observed when the anti-K^d^ phages were used alone, and mutations on the *cps* operon (*wcaJ*) were only observed in the NTUH_K2044/cp48+mtp6 experiment (as for NTUH_K2044/cp48+mtp5 and NTUH_K2044/mtp6).

### Both anti-K and anti-K^d^ phages can target wt *K. pneumoniae* within the mammalian gut

In order to investigate the ability of anti-K^d^ phages to infect Kp *in vivo, we* selected mtp5 and used it alone or in combination with phage cp1, against strain BJ1-GA (the original host of both phages) in mice gut colonization experiments. After a single dose of phage, a small non-statistically significant decrease of Kp abundance in the feces on the subsequent days was observed (**Figure S9, Table S6**). However, stable phage numbers were observed, pointing to replication of both phages mtp5 and cp1, when used either alone or in combination in the mice gut (**Figure S9**). When extending the phage administration for 2 more days, a decrease of the bacterial loads was observed again on day 4 the first day after treatment (**Figure 7AC**). Although a 10 to 100-fold decrease of the numbers of the host strain was observed in the feces, no statistical significance was obtained possibly due to high variability of the samples (**Figure 7C, Table S7**), we could again detect stable levels of phage and bacterial populations with three consecutive phage treatments (**Figure 7AB**). These observations indicate that infection and replication of both phages occurred in the mice gut.

**Figure 7.**
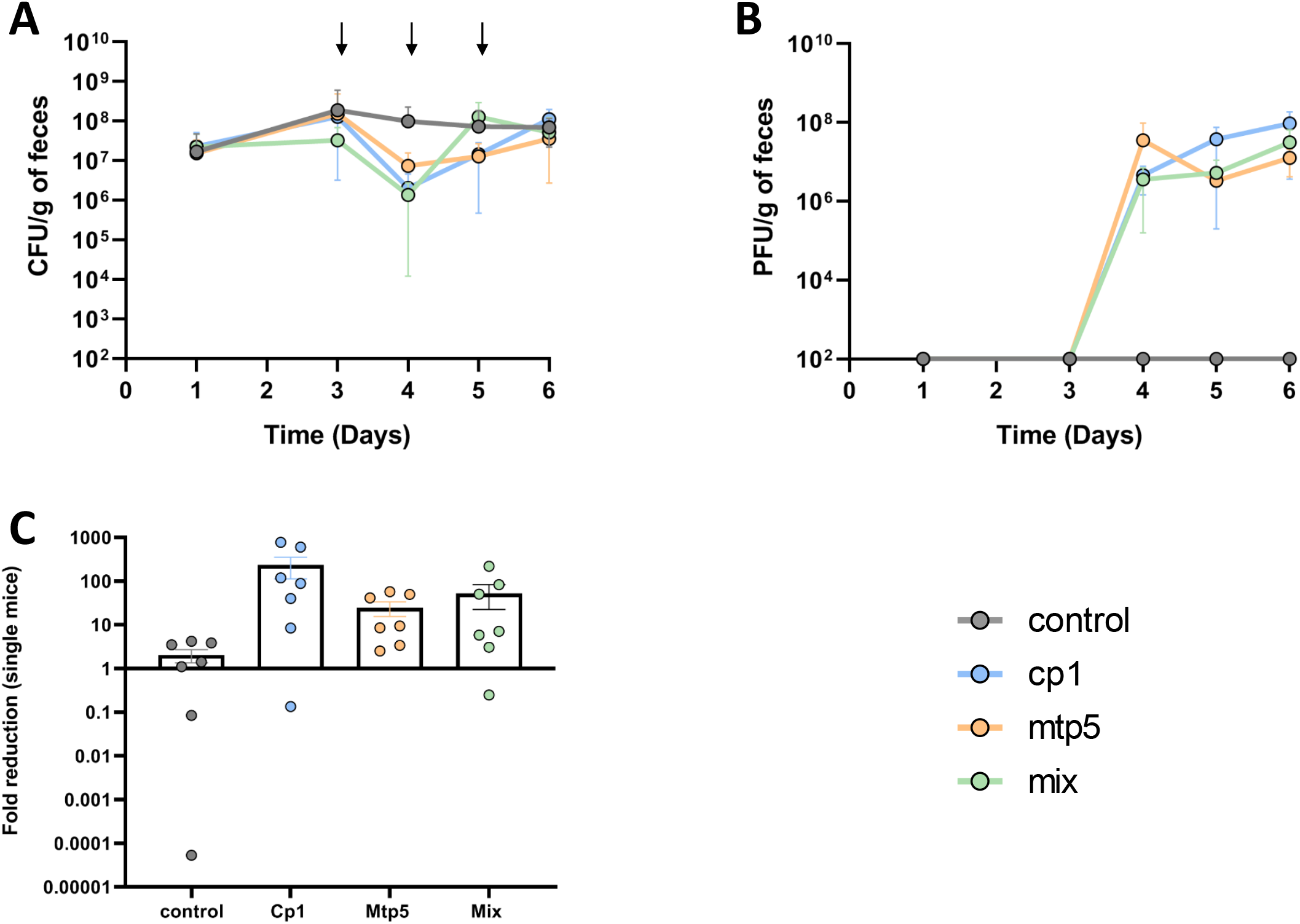
Phage replication and coexistence with *K. pneumoniae* in the mouse gut. *K. pneumoniae* BJ1GA-colonized OMM12 mice (n=28) received by oral gavage at day 3, 4 and 5 either PBS (pink, n=7) or phages cp1 and mtp5, together (mix; green, n=7; 6×10^7^ pfu per dose made of the same amount of each phage) or individually (cp1: blue, n=7; mtp5: orange, n=7). **A.** Levels of *K. pneumoniae* BJ1-GA in the feces (arrows represent the days at which phage(s) was/were given to the mice). **B.** Phage titers from the fecal samples reported in panel A. **C.** Fold-reduction of the bacterial population levels one day after first treatment.

We investigated whether phage-resistant mutants emerged *in vivo*, as previously observed *in vitro*. From the different fecal samples collected at days 4, 5 and 6, different colony phenotypes were observed: large colonies (possibly hypercapsulated), medium colonies and small colonies (**Figure S10**). In the control mice, from days 4 to 6, we only observed the presence of the large and medium colony phenotypes (**Figure S10_A**), and a similar pattern was observed for the mice treated with phage mtp5. In contrast, in the mice receiving phage cp1 and the cocktail (cp1+mtp5), the three different phenotypes were detected (**Figure S10_BCD**).

One colony of each phenotype was isolated per treatment and per day, and tested for resistance to both phages. For the control and mtp5 groups, both colony phenotypes showed susceptibility to phage cp1 (this was true for colonies recovered during the three days of phage administration), whereas for phage mtp5, resistant and intermediate resistant populations (as seen before, with lysis at high concentration but no plaques at lower dilutions) were observed at day 4 and at days 5 and 6 (**Figure S10_E**). On the other hand, the phenotypes isolated from mice treated with phage cp1 and the mtp5+cp1 cocktail showed variable susceptibility to the two phages (**Figure S10_E**). Surprisingly, we could detect colonies that were susceptible to both phages (d6/cp1-small and medium colonies and d5/mix-small colonies).

## Discussion

The structural diversity of capsular polysaccharides of *Klebsiella* is a major hurdle for the development of anti-*Klebsiella* phage therapy strategies: the capsule is normally abundant and thus acts as receptor for the majority of anti-*Klebsiella* phages described until now; hence, the phage host range is generally restricted to the K-type of the bacterial host from which the phage was isolated (Rieger-Hug and Stirm 1981; Lin et al. 2014; Fang et al. 2022; Eckstein et al. 2021; Chen et al. 2022; Fang and Zong 2022; Beamud et al. 2022). In some Kp infections, capsule-specialist phages may represent efficient tools, for example in the case of liver abscess, a localized infection which is mainly associated with K1 and K2 capsular types (Lin et al. 2014; Hung et al. 2011), but in most other infection sources, capsular diversity of Kp is very high.

Consistent with the literature, a marked host specificity was observed for anti-K phages isolated in this study, which were typically only able to target a unique KL-type. Exceptions were observed with a minority of these phages including cp8, cp10, cp11, cp12 and cp13, isolated from a KL10 host strain and also able to infect KL25 strains. This may be explained by the presence of different depolymerase domains in these phages (Table S4). Additionally, anti-K phages cp41, cp45 and cp39 infected KL106 wt strains but also OL1v1 capsule-deficient mutants. This may be explained by the fact that the widespread ST258-KL106 lineage produces capsule at low levels (Castronovo et al. 2017), and therefore the isolated phages may have non-capsular structures as receptors. This hypothesis is reinforced by the presence in these phages of a depolymerase domain also identified in some anti-K^d^ phages. Despite the dominant picture of capsule type restriction of anti-K phages, notable exceptions have also been described previously, with phages carrying multiple depolymerase domains. Still, the broadest-range phage was able to infect only 11 different KL-types (Pan et al. 2017; Šimoliūnas et al. 2013; Townsend et al. 2021), which remains limited in face of the approximately 200 K types described or inferred by Kp genomics.

*In vitro* studies have shown that capsulated Kp strains typically evolve phage resistance through capsule production inactivation (Hesse et al. 2020; D. Tan et al. 2020; Cai et al. 2019; Majkowska-Skrobek et al. 2021). Consistently, we found that emergence of resistance against anti-K phages is rapid and revolves mainly around mutations or disruptions mediated by ISs on the glycosyltransferases initiators of capsule production (*wcaJ /wbaP*) (Haudiquet et al. 2021)(Hesse et al. 2020; D. Tan et al. 2020; Cai et al. 2019). The disruption of *wbaP* due to insertion by an IS91 initially present in a IncF plasmid of the strain (NJST258/cp18 or cp20), corroborates the interplay between plasmids and phage pressure leading to facilitation of plasmid conjugation (Haudiquet et al. 2021).

In this study, we have followed an innovative approach to isolate phages targeting the more conserved structures that lie below the capsule (anti-K^d^ phages), using capsule-deficient mutants as hosts. We compare such phages for their genomic, *in vitro* and *in vivo* characteristics with anti-capsule phages, and explore the interactions between these two types of phages.

Consistent with our initial hypothesis, phylogenetically diverse anti-K^d^ phages were isolated. In addition, the anti-K^d^ phages infected a broad range of non-capsulated strains. In a recent study, phages showing a capsule-independent mode of entry also exhibited a much broader host range than phages against anti-capsulated strains (Beamud et al. 2022). The strongest illustration of a wide host range breadth of anti-K^d^ phages was provided by phage mtp5. This phage was able to target all tested non-capsulated strains, and to infect almost all wt Kp strains when combined with anti-K phages. For some of the strains infected with mtp5, we were not able to assess the efficiency of plating due to the absence of phage plaques on lower dilutions. This phenomenon was observed for strains belonging to different OL-types (OL1, OL2, OL3/O3a, OL4 and OL12), yet we were able to assess phage EOP on other strains with the same OL-types.

In addition to their broad host-range, anti-K^d^ phages showed improved resilience against the emergence of non-susceptible populations. Indeed, we observed extended time before regrowth of bacterial populations, and even more so when used in cocktail, in addition to a reduced relative bacterial growth, compared to anti-K phages. We also observed that depending on the anti-K^d^ phages, resistance is the result of a diversity of mutations, with the LPS core seemingly playing a prominent role in anti-K^d^ phages resistance, suggesting it may play a role as receptor. The conservation of this important membrane component is consistent with the broad host range exhibited by these phages. Still, resistance to mtp5, alone or in cocktail with anti-K phages, did not rely on mutation of a specific gene, operon or step of the LPS pathway, but rather disruption of several genes that play an important role in the process of membrane biosynthesis. The evolution by loss of particular membrane proteins or components of the core LPS, may delay the development of anti-K^d^ phage resistance by requesting specific mutations and by necessitating physiological adjustments against important changes to the cell membrane composition. Different from the capsule, the presence of which seems accessory or even deleterious in some environments, the loss of these conserved membrane components may incur a high fitness cost. Further work is necessary to test this hypothesis.

Whether capsule production is systematic, abundant and stable is essential for the prospect of phage therapy, as it impacts anti-K phage efficacy *in vivo*, and directly determines the relevance of an anti-K^d^ phage strategy. Our mouse colonization experiments showed replication of the anti-K phage, as expected, but also of the anti-K^d^ phage when used individually in the gut, with a small but transient reduction on the bacterial loads. These dynamics are in line with previous studies that suggested a coexistence of bacterial and virulent phage populations in the mice gut (Maura et al. 2012; Lourenço et al. 2020; B. B. Hsu et al. 2019; Mirzaei and Maurice 2017; Kirsch et al. 2021). This important observation suggests that capsule expression may not be systematic in the gut, either due to non-uniform expression along the gastrointestinal tract, or over time, or in specific sub-niches even in the absence of selection by phages (Lourenço et al. 2020; 2022). We observed the presence of three different colony phenotypes (**Figure S10**), and their phage susceptibility phenotypes corroborate the hypothesis that Kp variants that do not express capsule can co-exist in the gut with capsule-expressing ones. Non-capsulated variants would give access to the anti-K^d^ phage, with consequent infection and replication, as observed here for mtp5. The presence of the same resistance phenotypes in the control and mtp5 treated mice leads to the hypothesis that phage mtp5 takes advantage of the partial lack of *cps* production but does not play a major selective role on the overall population, while the cp1 phage plays a selective role due to the high representation of capsulated populations, leading to the emergence of different morphotypes and different resistance profiles.

The complexity of the gut environment leads to multiple ecological interactions, and capsule expression may indeed be associated with variable fitness costs (Y. H. Tan et al. 2020; Koskella et al. 2012). Different subpopulations of less capsulated or even entirely non-capsulated Kp may occupy different niches in the gut environment. Understanding the expression of Kp capsule in different sites *in vivo* will be important to define the potential of anti-K^d^ phages. The broad host-range observed for anti-K^d^ is coherent with these phages having access to small and possibly transient populations in the environment, leading to the selection of more generalist phages (Bono et al. 2015).

In summary, we show that anti-K^d^ phages, which we defined as anti-*Klebsiella* phages isolated from capsule-deficient strains, combine interesting properties for the prospect of phage therapy, including broad host range, slower emergence of resistance, and synergy with anti-K phages. Some anti-K^d^ phages uncovered in this work, such as mtp5, had a complete or nearly complete host-range (as assessed based on O-antigen diversity), suggesting that targeting conserved structures (possibly conserved more broadly beyond the Kp species itself) generally believed to be inaccessible due to the thick capsule, is a realistic avenue to circumvent the narrow spectrum conundrum of anti-K phages. Given its implications for anti-*Klebsiella* phage therapy and the potential role of anti-K^d^ phages, more work is needed on the important question of where, when and how much is the capsule expressed during colonization or infection.

## Material and Methods

### Bacterial strains and phages

Bacterial strains used for phage isolation and host-range assays are listed in **Table S2**. Strains were routinely cultured in lysogeny broth (LB), or on LB agar or Simmons Citrate agar with Inositol (SCAI) plates, at 37°C.

Phages isolated in this study are described in **Table S3**. Phages were amplified in exponential growing cultures of the respective host strain for approximately 4 to 5 hours (h). Cell lysate supernatants containing amplified phages were 0.22 μm-filter sterilized and stored at 4°C. Genome assemblies of the phages isolated in this study have been deposited in the European Nucleotide Archive under the BioProject PRJEB55170.

### Genomic background and surface structures analysis of publicly available genomes of *K. pneumoniae* species complex for phage host strains selection

The *K. pneumoniae* species complex includes five species distributed among seven phylogroups: *K. pneumoniae* subsp. *pneumoniae* (Kp1), *K. quasipneumoniae* subsp. *quasipneumoniae* (Kp2), *K. variicola* subsp. *variicola* (Kp3), *K. quasipneumoniae* subsp. *similipneumoniae* (Kp4), *K. variicola* subsp. *tropica* (Kp5), *‘K. quasivariicola’* (Kp6) and *K. africana* (Kp7) (Rodrigues et al. 2019). We have previously collated 7,388 *K. pneumoniae* genomic sequences from NCBI GenBank database on March 2019 (Hennart et al. 2021). Comparative genomic analysis were performed using Kleborate v1.0 (Lam, Wick, Watts, et al. 2021), a tool designed for the genotyping of *K. pneumoniae*. Kleborate can extract from the genomes relevant information such as species assignation, the classical 7-gene MLST sequence type (ST), several chromosomal and plasmid-associated virulence loci, antimicrobial resistance genes and mutations, and can predict (with a confidence score) the capsule type (KL) and O antigen locus (OL) type (Lam, Wick, Watts, et al. 2021).

Based on the genotyping information collected, we selected a total of 14 *K. pneumoniae* strains for phage isolation (**Table** 1). These included 7 wild-type (wt) strains (all Kp1) and 7 capsule-deficient mutants (6 Kp1 and 1 Kp3) (de Sousa et al. 2020), as hosts for isolation of anti-K phages and anti-K^d^ phages, respectively.

### Phage isolation and host range tests

Phage isolation was performed using water from the river Seine and from sewage, as previously described (Lourenço et al. 2020).

Host range tests were performed as follows: 3 μL of phosphate-buffered saline (PBS)-diluted phage solutions (0.22 μm filter sterilized crude lysates adjusted to 10^7^ PFU/mL) were deposited side by side on the lawn of each tested bacterium on square LB agar plates. Plates were incubated at 37°C overnight (≈18 h).

Isolated phages were tested against 50 different *K. pneumoniae* strains, including 43 wt strains representing 32 different KL-types and the 7 capsule-deficient mutants (de Sousa et al. 2020) (**Table S2**); and against 16 strains from other *Enterobacteriaceae* species (9 *Escherichia coli*, 6 *Salmonella* spp. and 1 *Citrobacter* spp.). Additionally, the anti-K^d^ phages were also tested against 23 *K. pneumoniae* non-mucoid (presumably capsule-deficient) clones generated as explained below (**Table S2**).

### Generation of non-mucoid *K. pneumoniae* strains (capsule-deficient clones)

Following Chiarelli *et al*. (Chiarelli et al. 2020), 23 wt *K. pneumoniae* strains were streaked on Tryptic Soy Agar (TSA) and incubated at 37°C for 24 h. Then, the TSA plates were scanned for non-mucoid (NM) sectors (**Figure S11**) at 25°C until NM sectors were observed. The sectors were then isolated, plated and kept at 37°C for another 24h, and then left at 25°C to verify if all colonies were homogenous. If so, two further steps of purification were performed before storage at −80°C (**Table S3**).

### Phage efficiency of plating (EOP) tests

The efficiency of plating (EOP) was calculated as the ratio of the number of plaques formed by the phage on each strain tested and the number of plaques formed on the specific host strain, for each phage. Three to four independent replicates were performed using bacterial cultures grown to an OD of approximately 0.2 at 600 nm and spread on LB plates onto which phage dilutions were spotted. Plates were incubated at 37°C overnight.

### Phage genome sequencing and analysis

Sterile phage lysates [obtained by a 5 h phage amplification, followed by centrifugation (10 min, 5000 rpm) and 0.22 μm filter sterilization] were treated with DNase (120 U) and RNase (240 mg/mL) for 30 min at 37°C before adding EDTA (20 mM). Lysates were then treated with proteinase K (100 mg/mL) and SDS (0.5%) at 55°C for 30 min. DNA was extracted by a phenol-chloroform protocol modified from Pickard 2009 (Pickard, 2009). Genomic DNA libraries were prepared with TruSeq DNA PCR-Free sample preparation kit (Illumina Inc., San Diego, USA) and 2Ͱ×Ͱ150 paired-end sequencing was performed using the NextSeq 500/550 Illumina technology (Illumina, San Diego, USA). The quality of the reads was checked with fastqc v0.8.5 (https://www.bioinformatics.babraham.ac.uk/projects/fastqc/) and reads were cleaned using the fqcleaner pipeline from Galaxy-Institut Pasteur (https://gitlab.pasteur.fr/GIPhy/fqCleanER). *De novo* assembly was performed using Spades (3.11.0) (Prjibelski et al. 2020), or using a workflow implemented in Galaxy-Institut Pasteur using clc_assembler v4.4.2 and clc_mapper v4.4.2 (CLC Bio, Qiagen) when necessary. Phage termini were determined by PhageTerm 2.0.1 (Garneau et al. 2017) and annotations performed using PATRIC RASTtk (Brettin et al. 2015; Aziz et al. 2008). Phage lifestyles were accessed using BACPHLIP, a python library for predicting phage lifestyle based on genome sequence (Hockenberry and Wilke 2021). The presence of genes coding for putative virulence factors or antibiotic resistance was investigated using Kleborate, Resfinder (antibiotic resistance), BigsDB Pasteur Klebsiella ‘Virulence genes’ and the VDFB virulence genes database (Bortolaia et al. 2020; Lam, Wick, Watts, et al. 2021; Liu et al. 2019; Bialek-Davenet et al. 2014).

Phylogenetic trees were generated using JolyTree v2.0 (MASH based; parameters: sketch size of 100000, probability of observing a random k-mer of 0.00001 and k-mer size of 15) (Criscuolo 2019), and visualized using iTol (Letunic and Bork 2019). 98 publicly available genomes deposited on the RefSeq NCBI database, with Kp (taxid: 573) stated as host (March 2021), were used for comparative analyses.

Average nucleotide identity (ANI) between similar phages was calculated using OGRI (https://gitlab.pasteur.fr/GIPhy/OGRI).

### Depolymerases analysis

To detect the putative depolymerases in our newly isolated phages, we performed an initial HMMER (v3.3) comparative analysis using 14 HMM profiles associated with bacteriophage-encoded depolymerases (de Sousa et al. 2020) (filtered by the e-value of the best domain maximum 10^−3^). A Blastp (v2.6.0, default parameters) filtering by e-value (maximum 10^−5^) and identity (30%) search was also performed for specific protein sequences described and validated for broad-range *Klebsiella* phages described in the literature (Pan et al. 2017) (**Table S4**).

### Growth curves and lysis kinetics analysis

To record phage replication and bacterial lysis, an overnight culture of the respective bacterial strain was diluted in LB Lennox media (Sigma-Aldrich) and grown to an OD at 600 nm of ≈0.2, from which 140 μL were distributed on a 96-well plate (Microtest 96 plates, Falcon). Afterwards were added 10 μL of sterile phage lysates, previously diluted in PBS, to obtain a multiplicity of infection (MOI) of 1 × 10^−2^ in each well. Plates were incubated in a microplate reader at 37°C, with a shaking step of 30 sec before the automatic recording of OD at 600 nm every 15 min over 18h-20 h (Tecan microplate reader). The Area Under the Curve (AUC) was calculated in R using the function trapz from the pracma package (v2.3.3). The Relative Bacterial Growth (RBG) was calculated using the following equation: RBG = [Abs600 (t = 4 h) -Abs600 (t = 0 h)]bp/[Abs600 (t = 4 h) - Abs600 (t = 0 h)]b, in which Abs600 stands for the absorbance at 600Ͱnm of bacterial cultures, b for bacteria only (control cultures), bp for bacteria with phage (infected cultures) and t for time in hours (h). RBG was calculated at 4, 6, 8 and 10 hours (Majkowska-Skrobek et al. 2021). Mann-Whitney-Wilcoxon test was performed for AUC and RBG results comparisons using R.

### Isolation and testing of putative phage-resistant clones

After 20 h of growth, cultures were plated in LB Lennox agar to select possible phage resistant clones. After ≈18 h at 37°C, colonies were counted and checked for phenotypic differences. In total, 24 colonies from the control cultures and 50 colonies from the infected cultures were isolated and grown in 96-well plates with LB Lennox plus 16% glycerol for 24 h at 37°C and stored at −80°C. Afterwards a fresh overnight culture of each clone (i.e. each isolated colony) was grown at 37°C and refreshed in LB liquid media the following day and grown at 37°C to an OD of approximately 0.5 (600 nm) and a double spot test (6 μl bacterial culture + 3 μl phage) was performed in order to test for resistance.

Subsequently, a fresh culture of 1 mL of each clone (*i.e*., each isolated colony) was grown for 24 h at 37°C and then all cultures were pooled together and centrifuged. This pelleted population was then diluted in 1 mL of PBS. 500 μL of the sample were used for DNA extraction using the Maxwell Cell tissue kit (Promega) and subjected to Illumina sequencing, as described above.

For some cultures, the detection of a very small colony variant phenotype led to difficulties in isolating them. In such cases, DNA extraction was performed directly from colonies collected from the plate, without a previous resistance test (**Table S9**).

Initial phenotypic observations (colonies on LB plates) regarding the anti-K phages showed an apparent loss of capsule within the population 20 h post-infection (to control for capsule presence, colonies were tested from samples in the presence and absence of phages). Colonies with the different phenotypes were counted before isolation for the resistance assay and subsequent DNA extraction and sequencing (**Table S8**).

### Animals and ethics

Oligo-MM12 C57BL/6NTac mice were bred in isolators (Getinge) in the germfree facility at the Helmholtz Center for Infection Research. Experiments were performed in gender and age matched animals, with female and male mice with an age of 8-12 weeks. Sterilized food and water was provided *ad libitum*. Mice were kept under strict 12-hour light cycles (lights on at 7:00 am and off at 7:00 pm) and housed in groups of 3-4 mice per cage. All mice were euthanized by asphyxiation with CO2 and cervical dislocation.

All animal experiments have been performed in agreement with the guidelines of the Helmholtz Center for Infection Research, Braunschweig, Germany; the National Animal Protection Law [Tierschutzgesetz (TierSchG) and Animal Experiment Regulations (Tierschutz-Versuchstierverordnung (TierSchVersV)], and the recommendations of the Federation of European Laboratory Animal Science Association (FELASA). The study was approved by the Lower Saxony State Office for Nature, Environment and Consumer Protection (LAVES), Oldenburg, Lower Saxony, Germany; permit No. 33.8-42502-04-20/3564.

### Murine model of *K. pneumoniae* colonization

*In vivo* colonization assays were performed using the OligoMM^12^ mice model. This mice model harbours a consortium of 12 different bacterial strains in the gut: *Acutalibacter muris, Akkermansia muciniphila, Bacteroides caecimuris, Bifidobacterium animalis, Blautia coccoides, Enterocloster clostridioformis, Clostridium innocuum, Enterococcus faecalis, Flavonifractor plautii, Limosilactobacillus reuteri, Muribaculum intestinale and Turicimonas muris* (Brugiroux et al. 2016). Two independent experiments with repeated phage administration were performed with n= 3-4 mice per group. At day 0, mice were orally colonized with Kp bacterial strain BJ1-GA (SB4496). Initial inoculum was prepared by culturing bacteria overnight at 37°C in LB broth with 25 mg/mL chloramphenicol. Subsequently, the culture was diluted 1:25 in fresh LB medium, and subcultured for 4 h at 37°C. Bacteria were resuspended in 10 mL phosphate-buffered saline (PBS) and adjusted to 5×10^8^ CFUs/ml. Mice were orally inoculated with 200 μL of bacterial suspension. At day 3 or day 3, 4 and 5 mice received the two phages alone (adjusted to 10^8^ PFU), or in combination (1:1 mixture of each phage in 200 μL of PBS) once daily by oral gavage.

The level of phages was assessed by serial dilutions in PBS and spotted on LB plates with an overlay of strain BJ1-GA. Weight of the mice was monitored during the course of the experiment, and feces were collected at different time points after colonization (day 1,3,4,5 and 6).

Subsequently, fecal samples were diluted in 1 mL PBS and homogenized by bead-beating with 1 mm zirconia/silica beads for two times 25 s using a Mini-Beadbeater-96 (BioSpec). To determine CFUs, serial dilutions of homogenized samples were plated on LB plates with 25 mg/mL chloramphenicol. To determine PFUs, serial dilutions of homogenized samples were plated on LB plates overlaid with strain BJ1-GA. Plates were incubated at 37°C over-night before counting. CFUs of strain BJ1-GA and PFUs of the phages were calculated after normalization to the weight of feces. For the isolation of clones with different phenotypes, feces samples of one mouse per group were plated on SCAI medium agar and one clone of each different phenotype was isolated in LB Lennox agar medium, followed by testing for susceptibility to phages cp1 and mtp5.

### Statistical Analysis

For the growth curves of the different strains exposed or not to phages (n=3 or n>3 for each condition), error bars represent standard error of the mean (SEM). To compare the Area under the curve (AUC) and relative bacterial growth (RBG) values, a Mann-Whitney test was performed (**P-value ≤ 0.01;***P-value ≤ 0.001).

Statistical analysis from *in vivo* mice experiments bacterial levels, were carried out using the lme4, lmerTest and car packages of R (Bates et al. 2014; Fox and Weisberg 2019; Kuznetsova, Brockhoff, and Christensen 2017). CFU numbers were log10-transformed prior to analysis. In each experiment, four groups of mice were considered, three groups exposed to the different phages (cp1, mtp5, mix) and an unexposed control group. The impact of phages was assessed based on the abundance of phages (log-PFU). Given the non-linearity of responses, the day at which a measure was performed was considered as a categorical variable. Linear mixed-models were used to account for random experimental effects (i.e., individuals and experiments effects).

Overall effects were assessed with Analysis of Variance (ANOVA) and post-hoc Tukey’s comparisons and were performed using the lsmeans R package (Lenth 2016). Only p< 0.05 was considered statistically significant.

## Authors contributions

SB and ML conceived the study. ML, VP and LO performed laboratory experiments. ML, FG and CR performed genomic analyses. SB and TS obtained funding. ML and SB drafted the paper. All authors contributed to and approved the submitted version.

## Acknowledgements

We thank Olaya Rendueles Garcia and Eduardo Rocha for sharing the mutant strains used for anti-K^d^ phage isolation. We thank the Biomics Platform, C2RT, Institut Pasteur, Paris, France, supported by France Génomique (ANR-10-INBS-09-09) and IBISA, specially Elodie Turc and Laure Lemée, for genomic libraries preparation and sequencing. We thank Melanie Hennart for bioinformatics methodological input. We are grateful to Jin-Town Wang for sharing strain NTUH-K2044. We thank Quentin Lamy-Besnier and Olaya Rendueles Garcia for critical reading of the manuscript.

## References

Abedon, Stephen T. 2011. ‘Lysis from Without’. Bacteriophage 1 (1): 46–49. https://doi.org/10.4161/bact.1.1.13980.

Aziz, Ramy K., Daniela Bartels, Aaron A. Best, Matthew DeJongh, Terrence Disz, Robert A. Edwards, Kevin Formsma, et al. 2008. ‘The RAST Server: Rapid Annotations Using Subsystems Technology’. BMC Genomics 9 (February): 75. https://doi.org/10.1186/1471-2164-9-75.

Bates, Douglas, Martin Mächler, Ben Bolker, and Steve Walker. 2014. ‘Fitting Linear Mixed-Effects Models Using Lme4’. ArXiv:1406.5823 [Stat], June. http://arxiv.org/abs/1406.5823.

Beamud, Beatriz, García-González, Mar Gómez-Ortega, Fernando González-Candelas, Pilar Domingo-Calap, and Rafael Sanjuan. 2022. ‘Genetic Determinants of Host Tropism in Klebsiella Phages’. BioRxiv, 1 June 2022. doi: https://doi.org/10.1101/2022.06.01.494021.

Bialek-Davenet, S., A. Criscuolo, F. Ailloud, V. Passet, L. Jones, A. S. Delannoy-Vieillard, B. Garin, et al. 2014. ‘Genomic Definition of Hypervirulent and Multidrug-Resistant Klebsiella Pneumoniae Clonal Groups’. Emerg Infect Dis 20 (11): 1812–20. https://doi.org/10.3201/eid2011.140206.

Bonilla, Estrada, Ana Rita Costa, Daan F. van den Berg, Teunke van Rossum, Stefan Hagedoorn, Hielke Walinga, Minfeng Xiao, et al. 2021. ‘Genomic Characterization of Four Novel Bacteriophages Infecting the Clinical Pathogen Klebsiella Pneumoniae’. DNA Research: An International Journal for Rapid Publication of Reports on Genes and Genomes 28 (4): dsab013. https://doi.org/10.1093/dnares/dsab013.

Bono, Lisa M., Catharine L. Gensel, David W. Pfennig, and Christina L. Burch. 2015. ‘Evolutionary Rescue and the Coexistence of Generalist and Specialist Competitors: An Experimental Test’. Proceedings of the Royal Society B: Biological Sciences 282 (1821): 20151932. https://doi.org/10.1098/rspb.2015.1932.

Bortolaia, Valeria, Rolf S. Kaas, Etienne Ruppe, Marilyn C. Roberts, Stefan Schwarz, Vincent Cattoir, Alain Philippon, et al. 2020. ‘ResFinder 4.0 for Predictions of Phenotypes from Genotypes’. The Journal of Antimicrobial Chemotherapy 75 (12): 3491–3500. https://doi.org/10.1093/jac/dkaa345.

Brettin, Thomas, James J. Davis, Terry Disz, Robert A. Edwards, Svetlana Gerdes, Gary J. Olsen, Robert Olson, et al. 2015. ‘RASTtk: A Modular and Extensible Implementation of the RAST Algorithm for Building Custom Annotation Pipelines and Annotating Batches of Genomes’. Scientific Reports 5 (February): 8365. https://doi.org/10.1038/srep08365.

Brugiroux, Sandrine, Markus Beutler, Carina Pfann, Debora Garzetti, Hans-Joachim Ruscheweyh, Diana Ring, Manuel Diehl, et al. 2016. ‘Genome-Guided Design of a Defined Mouse Microbiota That Confers Colonization Resistance against Salmonella Enterica Serovar Typhimurium’. Nature Microbiology 2 (November): 16215. https://doi.org/10.1038/nmicrobiol.2016.215.

Cai, Ruopeng, Gang Wang, Shuai Le, Mei Wu, Mengjun Cheng, Zhimin Guo, Yalu Ji, et al. 2019. ‘Three Capsular Polysaccharide Synthesis-Related Glucosyltransferases, GT-1, GT-2 and WcaJ, Are Associated With Virulence and Phage Sensitivity of Klebsiella Pneumoniae’. Frontiers in Microbiology 10: 1189. https://doi.org/10.3389/fmicb.2019.01189.

Castronovo, Giuseppe, Ann Maria Clemente, Alberto Antonelli, Marco Maria D’Andrea, Michele Tanturli, Eloisa Perissi, Sara Paccosi, et al. 2017. ‘Differences in Inflammatory Response Induced by Two Representatives of Clades of the Pandemic ST258 Klebsiella Pneumoniae Clonal Lineage Producing KPC-Type Carbapenemases’. PloS One 12 (1): e0170125. https://doi.org/10.1371/journal.pone.0170125.

Chen, Xinyi, Qi Tang, Xue Li, Xiangkuan Zheng, Pei Li, Min Li, Fei Wu, Zhengjun Xu, Renfei Lu, and Wei Zhang. 2022. ‘Isolation, Characterization, and Genome Analysis of Bacteriophage P929 That Could Specifically Lyase the KL19 Capsular Type of Klebsiella Pneumoniae’. Virus Research, March, 198750. https://doi.org/10.1016/j.virusres.2022.198750.

Chiarelli, Adriana, Nicolas Cabanel, Isabelle Rosinski-Chupin, Pengdbamba Dieudonné Zongo, Thierry Naas, Rémy A. Bonnin, and Philippe Glaser. 2020. ‘Diversity of Mucoid to Non-Mucoid Switch among Carbapenemase-Producing Klebsiella Pneumoniae’. BMC Microbiology 20 (1): 325. https://doi.org/10.1186/s12866-020-02007-y.

Criscuolo, Alexis. 2019. ‘A Fast Alignment-Free Bioinformatics Procedure to Infer Accurate Distance-Based Phylogenetic Trees from Genome Assemblies’. Research Ideas and Outcomes 5 (June): e36178. https://doi.org/10.3897/rio.5.e36178.

De Sordi, Luisa, Varun Khanna, and Laurent Debarbieux. 2017. ‘The Gut Microbiota Facilitates Drifts in the Genetic Diversity and Infectivity of Bacterial Viruses’. Cell Host & Microbe 22 (6): 801–808.e3. https://doi.org/10.1016/j.chom.2017.10.010.

Eckstein, Simone, Jana Stender, Sonia Mzoughi, Kilian Vogele, Jana Kühn, Daniela Friese, Christina Bugert, et al. 2021. ‘Isolation and Characterization of Lytic Phage TUN1 Specific for Klebsiella Pneumoniae K64 Clinical Isolates from Tunisia’. BMC Microbiology 21 (1): 186. https://doi.org/10.1186/s12866-021-02251-w.

Ernst, Christoph M., Julian R. Braxton, Carlos A. Rodriguez-Osorio, Anna P. Zagieboylo, Li Li, Alejandro Pironti, Abigail L. Manson, et al. 2020. ‘Adaptive Evolution of Virulence and Persistence in Carbapenem-Resistant Klebsiella Pneumoniae’. Nature Medicine 26 (5): 705–11. https://doi.org/10.1038/s41591-020-0825-4.

Fang, Qingqing, Yu Feng, Alan McNally, and Zhiyong Zong. 2022. ‘Characterization of Phage Resistance and Phages Capable of Intestinal Decolonization of Carbapenem-Resistant Klebsiella Pneumoniae in Mice’. Communications Biology 5 (1): 48. https://doi.org/10.1038/s42003-022-03001-y.

Fang, Qingqing, and Zhiyong Zong. 2022. ‘Lytic Phages against ST11 K47 Carbapenem-Resistant Klebsiella Pneumoniae and the Corresponding Phage Resistance Mechanisms’. MSphere, March, e0008022. https://doi.org/10.1128/msphere.00080-22.

Follador, Rainer, Eva Heinz, Kelly L. Wyres, Matthew J. Ellington, Michael Kowarik, Kathryn E. Holt, and Nicholas R. Thomson. 2016. ‘The Diversity of Klebsiella Pneumoniae Surface Polysaccharides’. Microbial Genomics 2 (8): e000073. https://doi.org/10.1099/mgen.0.000073.

Fox, John, and Sanford Weisberg. 2019. An R Companion to Applied Regression. Third edition. Los Angeles: SAGE.

Garneau, Julian R., Florence Depardieu, Louis-Charles Fortier, David Bikard, and Marc Monot. 2017. ‘PhageTerm: A Tool for Fast and Accurate Determination of Phage Termini and Packaging Mechanism Using next-Generation Sequencing Data’. Scientific Reports 7 (1): 8292. https://doi.org/10.1038/s41598-017-07910-5.

Gorodnichev, Roman B., Nikolay V. Volozhantsev, Valentina M. Krasilnikova, Ivan N. Bodoev, Maria A. Kornienko, Nikita S. Kuptsov, Anastasia V. Popova, et al. 2021. ‘Novel Klebsiella Pneumoniae K23-Specific Bacteriophages From Different Families: Similarity of Depolymerases and Their Therapeutic Potential’. Frontiers in Microbiology 12: 669618. https://doi.org/10.3389/fmicb.2021.669618.

Hao, Guijuan, Rundong Shu, Liqin Ding, Xia Chen, Yonghao Miao, Jiaqi Wu, Haijian Zhou, and Hui Wang. 2021. ‘Bacteriophage SRD2021 Recognizing Capsular Polysaccharide Shows Therapeutic Potential in Serotype K47 Klebsiella Pneumoniae Infections’. Antibiotics (Basel, Switzerland) 10 (8): 894. https://doi.org/10.3390/antibiotics10080894.

Haudiquet, Matthieu, Amandine Buffet, Olaya Rendueles, and Eduardo P. C. Rocha. 2021. ‘Interplay between the Cell Envelope and Mobile Genetic Elements Shapes Gene Flow in Populations of the Nosocomial Pathogen Klebsiella Pneumoniae’. PLoS Biology 19 (7): e3001276. https://doi.org/10.1371/journal.pbio.3001276.

Hennart, Melanie, Julien Guglielmini, Martin C.J. Maiden, Keith A. Jolley, Alexis Criscuolo, and Sylvain Brisse. 2021. ‘A Dual Barcoding Approach to Bacterial Strain Nomenclature: Genomic Taxonomy of *Klebsiella Pneumoniae* Strains’. Preprint. Microbiology. https://doi.org/10.1101/2021.07.26.453808.

Hernandez, Catherine A., and Britt Koskella. 2019. ‘Phage Resistance Evolution in Vitro Is Not Reflective of in Vivo Outcome in a Plant-Bacteria-Phage System’. Evolution; International Journal of Organic Evolution 73 (12): 2461–75. https://doi.org/10.1111/evo.13833.

Hesse, Shayla, Manoj Rajaure, Erin Wall, Joy Johnson, Valery Bliskovsky, Susan Gottesman, and Sankar Adhya. 2020. ‘Phage Resistance in Multidrug-Resistant Klebsiella Pneumoniae ST258 Evolves via Diverse Mutations That Culminate in Impaired Adsorption’. MBio 11 (1): e02530–19. https://doi.org/10.1128/mBio.02530-19.

Hockenberry, Adam J., and Claus O. Wilke. 2021. ‘BACPHLIP: Predicting Bacteriophage Lifestyle from Conserved Protein Domains’. PeerJ 9: e11396. https://doi.org/10.7717/peerj.11396.

Hoyles, Lesley, James Murphy, Horst Neve, Knut J. Heller, Jane F. Turton, Jennifer Mahony, Jeremy D. Sanderson, et al. 2015. ‘Klebsiella Pneumoniae Subsp. Pneumoniae-Bacteriophage Combination from the Caecal Effluent of a Healthy Woman’. PeerJ 3: e1061. https://doi.org/10.7717/peerj.1061.

Hsu, Bryan B., Travis E. Gibson, Vladimir Yeliseyev, Qing Liu, Lorena Lyon, Lynn Bry, Pamela A. Silver, and Georg K. Gerber. 2019. ‘Dynamic Modulation of the Gut Microbiota and Metabolome by Bacteriophages in a Mouse Model’. Cell Host & Microbe 25 (6): 803–814.e5. https://doi.org/10.1016/j.chom.2019.05.001.

Hsu, C. R., T. L. Lin, Y. C. Chen, H. C. Chou, and J. T. Wang. 2011. ‘The Role of Klebsiella Pneumoniae RmpA in Capsular Polysaccharide Synthesis and Virulence Revisited’. Microbiology 157 (Pt 12): 3446–57. https://doi.org/10.1099/mic.0.050336-0.

Hung, Chih-Hsin, Chih-Feng Kuo, Chiou-Huey Wang, Ching-Ming Wu, and Nina Tsao. 2011. ‘Experimental Phage Therapy in Treating Klebsiella Pneumoniae-Mediated Liver Abscesses and Bacteremia in Mice’. Antimicrobial Agents and Chemotherapy 55 (4): 1358–65. https://doi.org/10.1128/AAC.01123-10.

Keseler, Ingrid M., Julio Collado-Vides, Alberto Santos-Zavaleta, Martin Peralta-Gil, Socorro Gama-Castro, Luis Muñiz-Rascado, César Bonavides-Martinez, et al. 2011. ‘EcoCyc: A Comprehensive Database of Escherichia Coli Biology’. Nucleic Acids Research 39 (Database issue): D583–590. https://doi.org/10.1093/nar/gkq1143.

Kirsch, Joshua M., Robert S. Brzozowski, Dominick Faith, June L. Round, Patrick R. Secor, and Breck A. Duerkop. 2021. ‘Bacteriophage-Bacteria Interactions in the Gut: From Invertebrates to Mammals’. Annual Review of Virology 8 (1): 95–113. https://doi.org/10.1146/annurev-virology-091919-101238.

Koberg, Sabrina, Erik Brinks, Gregor Fiedler, Christina Hüsing, Gyu-Sung Cho, Marc P. Hoeppner, Knut J. Heller, Horst Neve, and Charles M. A. P. Franz. 2017. ‘Genome Sequence of Klebsiella Pneumoniae Bacteriophage PMBT1 Isolated from Raw Sewage’. Genome Announcements 5 (8): e00914–16. https://doi.org/10.1128/genomeA.00914-16.

Koskella, Britt, Derek M. Lin, Angus Buckling, and John N. Thompson. 2012. ‘The Costs of Evolving Resistance in Heterogeneous Parasite Environments’. Proceedings. Biological Sciences 279 (1735): 1896–1903. https://doi.org/10.1098/rspb.2011.2259.

Kuznetsova, Alexandra, Per B. Brockhoff, and Rune H. B. Christensen. 2017. ‘**LmerTest** Package: Tests in Linear Mixed Effects Models’. Journal of Statistical Software 82 (13). https://doi.org/10.18637/jss.v082.i13.

Lam, Margaret M. C., Ryan R. Wick, Louise M. Judd, Kathryn E. Holt, and Kelly L. Wyres. 2021. ‘Kaptive 2.0: Updated Capsule and LPS Locus Typing for the *Klebsiella Pneumoniae* Species Complex’. Preprint. Genomics. https://doi.org/10.1101/2021.11.05.467534.

Lam, Margaret M. C., Ryan R. Wick, Stephen C. Watts, Louise T. Cerdeira, Kelly L. Wyres, and Kathryn E. Holt. 2021. ‘A Genomic Surveillance Framework and Genotyping Tool for Klebsiella Pneumoniae and Its Related Species Complex’. Nature Communications 12 (1): 4188. https://doi.org/10.1038/s41467-021-24448-3.

Lenth, Russell V. 2016. ‘Least-Squares Means: The R Package Lsmeans’. Journal of Statistical Software 69 (1). https://doi.org/10.18637/jss.v069.i01.

Letunic, Ivica, and Peer Bork. 2019. ‘Interactive Tree Of Life (ITOL) v4: Recent Updates and New Developments’. Nucleic Acids Research 47 (W1): W256–59. https://doi.org/10.1093/nar/gkz239.

Lin, Tzu-Lung, Pei-Fang Hsieh, Yu-Tsung Huang, Wei-Ching Lee, Yi-Ting Tsai, Po-An Su, Yi-Jiun Pan, Chun-Ru Hsu, Meng-Chuan Wu, and Jin-Town Wang. 2014. ‘Isolation of a Bacteriophage and Its Depolymerase Specific for K1 Capsule of Klebsiella Pneumoniae: Implication in Typing and Treatment’. The Journal of Infectious Diseases 210 (11): 1734–44. https://doi.org/10.1093/infdis/jiu332.

Liu, Bo, Dandan Zheng, Qi Jin, Lihong Chen, and Jian Yang. 2019. ‘VFDB 2019: A Comparative Pathogenomic Platform with an Interactive Web Interface’. Nucleic Acids Research 47 (D1): D687–92. https://doi.org/10.1093/nar/gky1080.

Lourenço, Marta, Lorenzo Chaffringeon, Quentin Lamy-Besnier, Thierry Pédron, Pascal Campagne, Claudia Eberl, Marion Bérard, Bärbel Stecher, Laurent Debarbieux, and Luisa De Sordi. 2020. ‘The Spatial Heterogeneity of the Gut Limits Predation and Fosters Coexistence of Bacteria and Bacteriophages’. Cell Host & Microbe 28 (3): 390–401.e5. https://doi.org/10.1016/j.chom.2020.06.002.

Lourenço, Marta, Lorenzo Chaffringeon, Quentin Lamy-Besnier, Marie Titécat, Thierry Pédron, Odile Sismeiro, Rachel Legendre, et al. 2022. ‘The Gut Environment Regulates Bacterial Gene Expression Which Modulates Susceptibility to Bacteriophage Infection’. Cell Host & Microbe 30 (4): 556–569.e5. https://doi.org/10.1016/j.chom.2022.03.014.

Maciejewska, Barbara, Bartosz Roszniowski, Akbar Espaillat, Agata Kęsik-Szeloch, Grazyna Majkowska-Skrobek, Andrew M. Kropinski, Yves Briers, Felipe Cava, Rob Lavigne, and Zuzanna Drulis-Kawa. 2017. ‘Klebsiella Phages Representing a Novel Clade of Viruses with an Unknown DNA Modification and Biotechnologically Interesting Enzymes’. Applied Microbiology and Biotechnology 101 (2): 673–84. https://doi.org/10.1007/s00253-016-7928-3.

Majkowska-Skrobek, Grazyna, Pawel Markwitz, Ewelina Sosnowska, Cédric Lood, Rob Lavigne, and Zuzanna Drulis-Kawa. 2021. ‘The Evolutionary Trade-Offs in Phage-Resistant Klebsiella Pneumoniae Entail Cross-Phage Sensitization and Loss of Multidrug Resistance’. Environmental Microbiology 23 (12): 7723–40. https://doi.org/10.1111/1462-2920.15476.

Maura, Damien, Eric Morello, Laurence du Merle, Perrine Bomme, Chantal Le Bouguénec, and Laurent Debarbieux. 2012. ‘Intestinal Colonization by Enteroaggregative Escherichia Coli Supports Long-Term Bacteriophage Replication in Mice’. Environmental Microbiology 14 (8): 1844–54. https://doi.org/10.1111/j.1462-2920.2011.02644.x.

Mavrich, Travis N., and Graham F. Hatfull. 2017. ‘Bacteriophage Evolution Differs by Host, Lifestyle and Genome’. Nature Microbiology 2 (July): 17112. https://doi.org/10.1038/nmicrobiol.2017.112.

Mijalis, Eleni M., Lauren E. Lessor, Jesse L. Cahill, Eric S. Rasche, and Gabriel F. Kuty Everett. 2015. ‘Complete Genome Sequence of Klebsiella Pneumoniae Carbapenemase-Producing K. Pneumoniae Myophage Miro’. Genome Announcements 3 (5): e01137–15. https://doi.org/10.1128/genomeA.01137-15.

Mirzaei, Mohammadali Khan, and Corinne F. Maurice. 2017. ‘Ménage à Trois in the Human Gut: Interactions between Host, Bacteria and Phages’. Nature Reviews. Microbiology 15 (7): 397–408. https://doi.org/10.1038/nrmicro.2017.30.

Ørskov, I., and F. Ørskov. 1984. ‘4 Serotyping of Klebsiella’. In Methods in Microbiology, 14:143–64. Elsevier. https://doi.org/10.1016/S0580-9517(08)70449-5.

Pan, Yi-Jiun, Tzu-Lung Lin, Ching-Ching Chen, Yun-Ting Tsai, Yi-Hsiang Cheng, Yi-Yin Chen, Pei-Fang Hsieh, Yi-Tsung Lin, and Jin-Town Wang. 2017. ‘Klebsiella Phage ΦK64-1 Encodes Multiple Depolymerases for Multiple Host Capsular Types’. Journal of Virology 91 (6): e02457–16. https://doi.org/10.1128/JVI.02457-16.

Pertics, Botond Zsombor, Alysia Cox, Adrienn Nyúl, Nóra Szamek, Tamás Kovács, and György Schneider. 2021. ‘Isolation and Characterization of a Novel Lytic Bacteriophage against the K2 Capsule-Expressing Hypervirulent Klebsiella Pneumoniae Strain 52145, and Identification of Its Functional Depolymerase’. Microorganisms 9 (3): 650. https://doi.org/10.3390/microorganisms9030650.

Prjibelski, Andrey, Dmitry Antipov, Dmitry Meleshko, Alla Lapidus, and Anton Korobeynikov. 2020. ‘Using SPAdes De Novo Assembler’. Current Protocols in Bioinformatics 70 (1): e102. https://doi.org/10.1002/cpbi.102.

Provasek, Vincent E., Lauren E. Lessor, Jesse L. Cahill, Eric S. Rasche, and Gabriel F. Kuty Everett. 2015. ‘Complete Genome Sequence of Carbapenemase-Producing Klebsiella Pneumoniae Myophage Matisse’. Genome Announcements 3 (5): e01136–15. https://doi.org/10.1128/genomeA.01136-15.

Rawlings, M., and J. E. Cronan. 1992. ‘The Gene Encoding Escherichia Coli Acyl Carrier Protein Lies within a Cluster of Fatty Acid Biosynthetic Genes’. The Journal of Biological Chemistry 267 (9): 5751–54.

Rieger-Hug, D., and S. Stirm. 1981. ‘Comparative Study of Host Capsule Depolymerases Associated with Klebsiella Bacteriophages’. Virology 113 (1): 363–78. https://doi.org/10.1016/0042-6822(81)90162-8.

Roach, Dwayne R., and Laurent Debarbieux. 2017. ‘Phage Therapy: Awakening a Sleeping Giant’. Emerging Topics in Life Sciences 1 (1): 93–103. https://doi.org/10.1042/ETLS20170002.

Rodrigues, Carla, Siddhi Desai, Virginie Passet, Devarshi Gajjar, and Sylvain Brisse. 2022. ‘Genomic Evolution of the Globally Disseminated Multidrug-Resistant Klebsiella Pneumoniae Clonal Group 147’. Microbial Genomics 8 (1). https://doi.org/10.1099/mgen.0.000737.

Rodrigues, Carla, Virginie Passet, Andriniaina Rakotondrasoa, Thierno Abdoulaye Diallo, Alexis Criscuolo, and Sylvain Brisse. 2019. ‘Description of Klebsiella Africanensis Sp. Nov., Klebsiella Variicola Subsp. Tropicalensis Subsp. Nov. and Klebsiella Variicola Subsp. Variicola Subsp. Nov.’ Research in Microbiology 170 (3): 165–70. https://doi.org/10.1016/j.resmic.2019.02.003.

Rodrigues, Carla, Clara Sousa, João A. Lopes, Ângela Novais, and Luísa Peixe. 2020. ‘A Front Line on Klebsiella Pneumoniae Capsular Polysaccharide Knowledge: Fourier Transform Infrared Spectroscopy as an Accurate and Fast Typing Tool’. MSystems 5 (2): e00386–19. https://doi.org/10.1128/mSystems.00386-19.

Šimoliunas, Eugenijus, Laura Kaliniene, Lidija Truncaitė, Aurelija Zajančkauskaitė, Juozas Staniulis, Algirdas Kaupinis, Marija Ger, Mindaugas Valius, and Rolandas Meškys. 2013. ‘Klebsiella Phage VB_KleM-RaK2 — A Giant Singleton Virus of the Family Myoviridae’. Edited by Mark J. van Raaij. PLoS ONE 8 (4): e60717. https://doi.org/10.1371/journal.pone.0060717.

Sousa, Jorge A. M. de, Amandine Buffet, Matthieu Haudiquet, Eduardo P. C. Rocha, and Olaya Rendueles. 2020. ‘Modular Prophage Interactions Driven by Capsule Serotype Select for Capsule Loss under Phage Predation’. The ISME Journal 14 (12): 2980–96. https://doi.org/10.1038/s41396-020-0726-z.

Tan, Demeng, Yiyuan Zhang, Jinhong Qin, Shuai Le, Jingmin Gu, Li-Kuang Chen, Xiaokui Guo, and Tongyu Zhu. 2020. ‘A Frameshift Mutation in WcaJ Associated with Phage Resistance in Klebsiella Pneumoniae’. Microorganisms 8 (3): E378. https://doi.org/10.3390/microorganisms8030378.

Tan, Yi Han, Yahua Chen, Wilson H. W. Chu, Lok-To Sham, and Yunn-Hwen Gan. 2020. ‘Cell Envelope Defects of Different Capsule-Null Mutants in K1 Hypervirulent Klebsiella Pneumoniae Can Affect Bacterial Pathogenesis’. Molecular Microbiology 113 (5): 889–905. https://doi.org/10.1111/mmi.14447.

Townsend, Eleanor M., Lucy Kelly, Lucy Gannon, George Muscatt, Rhys Dunstan, Slawomir Michniewski, Hari Sapkota, et al. 2021. ‘Isolation and Characterization of Klebsiella Phages for Phage Therapy’. PHAGE (New Rochelle, N.Y.) 2 (1): 26–42. https://doi.org/10.1089/phage.2020.0046.

Wick, Ryan R., Eva Heinz, Kathryn E. Holt, and Kelly L. Wyres. 2018. ‘Kaptive Web: User-Friendly Capsule and Lipopolysaccharide Serotype Prediction for Klebsiella Genomes’. Journal of Clinical Microbiology 56 (6): e00197–18. https://doi.org/10.1128/JCM.00197-18.

Wyres, Kelly L., Ryan R. Wick, Louise M. Judd, Roni Froumine, Alex Tokolyi, Claire L. Gorrie, Margaret M. C. Lam, Sebastián Duchêne, Adam Jenney, and Kathryn E. Holt. 2019. ‘Distinct Evolutionary Dynamics of Horizontal Gene Transfer in Drug Resistant and Virulent Clones of Klebsiella Pneumoniae’. PLoS Genetics 15 (4): e1008114. https://doi.org/10.1371/journal.pgen.1008114.

