## supplementary material for "Phages against non-capsulated *Klebsiella pneumoniae*: broader host range, slower resistance"

**Supplementary text:**

**1 - Strain selection for phage isolation**

Based on the population genomics analysis (**Figure S2**), we selected 7 *K. pneumoniae* (sensu stricto) strains as hosts for anti-capsulated strain phage isolation:

- *K. pneumoniae* NJST258_2 (SB4975), a representative of the ST258-KL107, one of the most widespread lineages in clinical settings and linked to nosocomial infections and outbreaks (Deleo et al. 2014)
- *K. pneumoniae* NCTC8172 (SB504), a ST505 harbouring the *cps* locus (KL) structure type 64, one of the most disseminated KL-types and which has been extensively associated with two widespread carbapenemase-producing ST's, ST147 and ST11 (Rodrigues et al. 2022; Dong et al. 2018).
- Two other selected hosts were strains SB5442 and SB5521, representing two successful MDR ST-KL combinations in clinical settings, ST101-KL106 and ST307-KL102, respectively (Huynh et al. 2020; Wyres et al. 2019).
- *K. pneumoniae* strain CIP52.214 (SB3245), a ST297-KL10, was selected as it represented one of the most prevalent KL-types.
- To finish, we selected two hypervirulent (Hv) *K. pneumoniae* strains, SB3341 ST66-KL2 and NTUH_K2044 (SB3928) ST23-KL1, representing K1 and K2 capsule types associated with severe invasive infections (Wu et al. 2009; Lery et al. 2014). ST23 is the most frequent sublineage isolated in community-acquired liver abscesses, which is a prevalent infection in Asian countries (Wu et al. 2009; Turton et al. 2007; Merlet et al. 2012).

We next selected seven capsule-deficient strains to be used for anti-$K^d$ phage isolation. These included 6 *K. pneumoniae* and 1 *K. variicola* subsp. *variicola* (Kp3): Kp1 SB20*Δwza* ST15/O1v1 (04A025), Kp1 SB3928*Δwza* ST23/O1v2 (NTUH_K2044), Kp1 SB4021*ΔwcaJ* (SA1) and SB4454*ΔwcaJ* (CG43) ST86/O1v1; Kp1 SB4496*Δwza* ST380/O1v1 (BJ1-GA), Kp1 SB4975*Δwza* ST258/O2v2 (NJST258_2) and

Kp3 SB579*Δwza* ST146-O3/O3a (342) (de Sousa et al. 2020). Together, the O types of these strains represented 50.7% of non-redundant strains in the genomic database.

**2- Anti-K phage cp48 and anti-K$^d$ phage mtp6 may share the same receptor**

Anti-K phage cp48 and anti-K$^d$ phage mtp6 infected the wt strain and the non-capsulated NTUH-K2044 mutant, respectively and exclusively. Comparison of these two phages revealed a high genomic similarity (99.8%), with the same tail fibers genes (**Figure S12**) suggesting that cp48 and mtp6 have the same receptor despite having been isolated from capsulated and capsule-deficient strains, respectively. Only 77 bp differences, distributed across three different regions, were detected between these two phages. First, on protein 18 from phage mtp6 (protein 47 in cp48), we observed several SNPs leading to a 7 amino acid (aa) change in the middle region of the protein. This protein was annotated as “phage protein” and when searched for similarity on NCBI database it was closely related to other hypothetical proteins. The second difference is an insertion of 77 bp on an HNH endonuclease Gp2.8/Gp7.7 of phage cp48 disrupting the protein in two parts: protein 21 of phage mtp6, divided into proteins 43 and 44 on phage cp48. HNH endonucleases have been shown to be associated with the terminases of a large number of diverse phages (Kala et al. 2014). Third, a SNP on protein 62 of phage mtp6 (protein 2 in cp48) at position 781 of the gene, lead to the presence of an aspartic acid on position 261 in the protein of mtp6 phage instead of asparagine on cp48. Comparison analysis with the NCBI protein database showed high similarity of the mtp6 CDS 62/ cp48 CDS 2 to another phage particle associated lyase. Phage lyases are known to degrade specific membrane polysaccharides, which in this case may be the cause on the differential infection affinity observed towards the capsulated and non-capsulated strains. Resistance to the cocktail of cp48 and mtp6 phages was conferred by an interruption of the wcaJ gene by an IS5 (**Figure S12**), as previously observed when this strain and its capsule-deficient mutant were separately infected with cp48 and mtp6, respectively.

**Supplementary tables:**

*Please see the separate Excel file for access to the supplementary tables.*

**Table S1.** Dataset of 7,388 K. pneumoniae species complex public genomes

**Table S2**. Collection of bacterial strains used in this study

**Table S3.** Main characteristics of the phages isolated in this study

**Table S4.** Depolymerases analysis

**Table S5.** Analysis of the acquired mutations after phage infection in vitro (Breseq analysis)

**Table S6.** Statistics - 1 day phage treatment

**Table S7.** Statistics - 3 days phage treatment

**Table S8**. Percentage of resistant clones *in vivo*

**Supplementary figures:**

**Figure S1**. Frequency of *Klebsiella pneumoniae* genomes breakdown by ST, KL or OL characteristics.

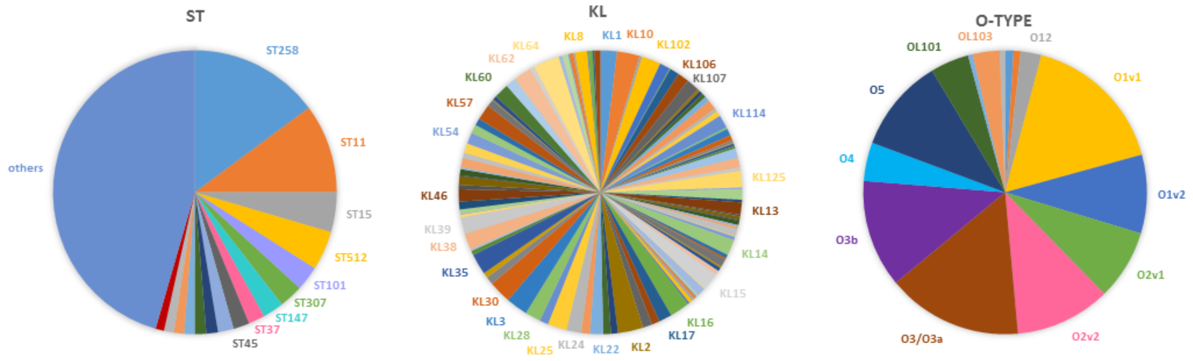


The ST distribution analysis was performed based on a dataset of 7388 genomes. To eliminate bias due to recent clonal expansions or outbreaks, KL- and O-antigen types frequencies were calculated using a non-redundant dataset of 1193 genomes, in which only one random strain per ST was included.

Based on the MLST, the seven most prevalent allelic profiles present in the dataset (7,388 genomes) were ST258, ST11, ST15, ST512, ST101, ST307 and ST147. Analysis of the subset showed KL64, KL2, KL10, KL30, KL3, KL35, KL16 and KL102 to be the most prevalent capsular types, and the most frequent O-antigen types were O1v1, O3/O3a, O3b, O5 and O2v2.

**Figure S2. Host range of anti-K phages**

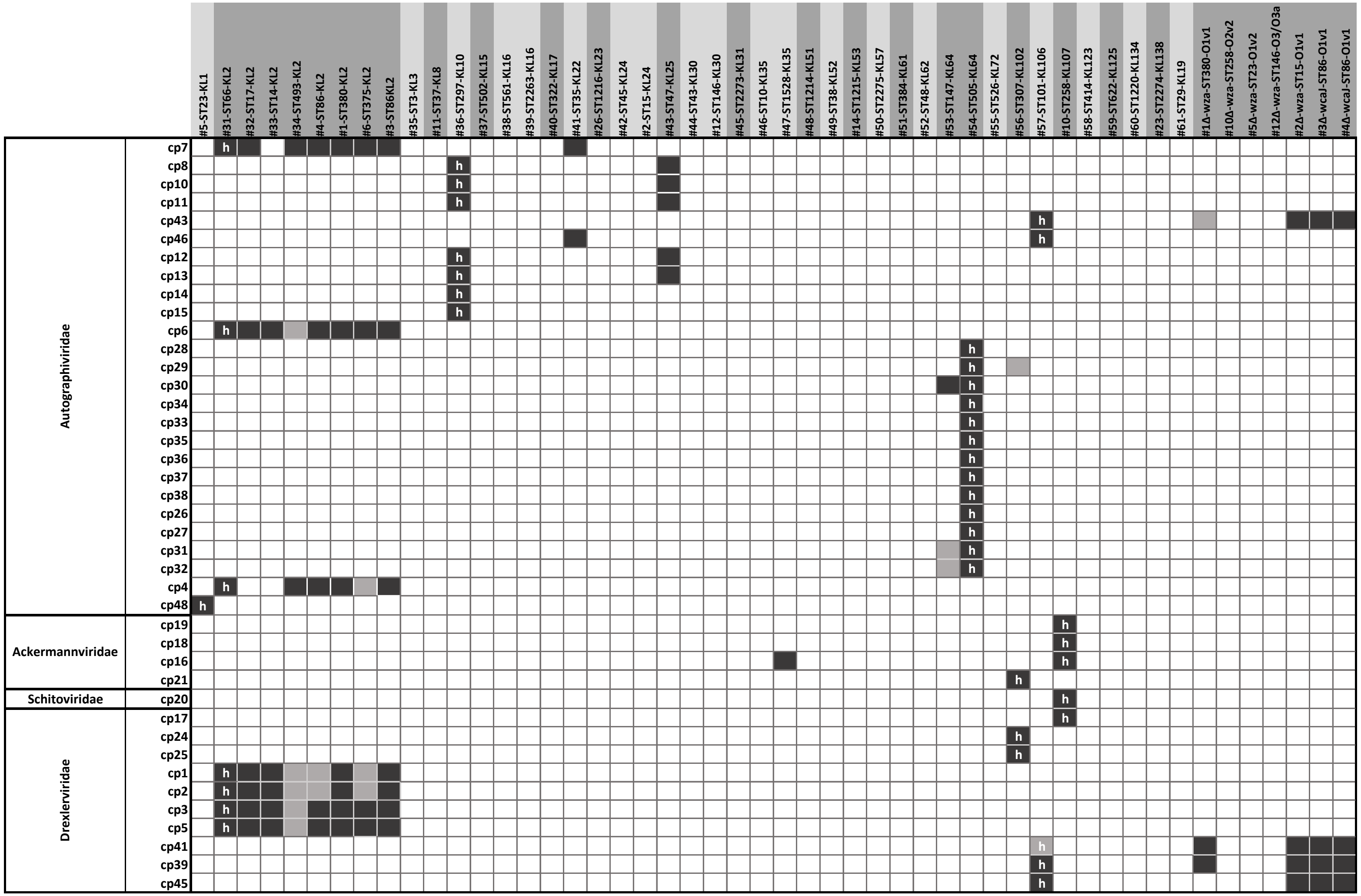

Anti-K phages were tested against wild-type strains and the 7 capsule-deficient (Δwza or ΔwcaJ) mutants. The dark-filled cells represent complete lysis, light-grey cells represent intermediate lysis and empty cells represent absence of lysis. “h” indicates the strain used as host for phage isolation.

**Figure S3.** Lysis kinetics of phage mtp5 on the 7 capsule-deficient mutant (Δ*wza* or Δ*wcaJ*) strains

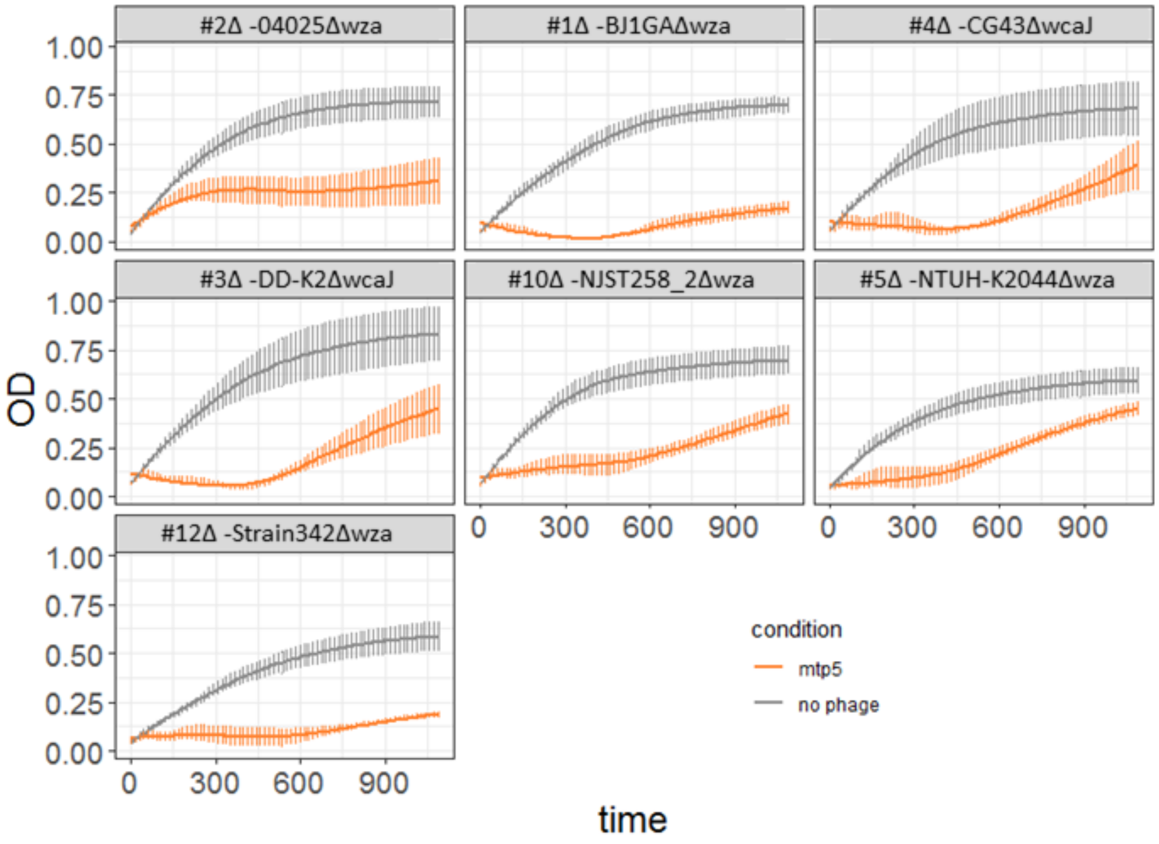


Growth curves of the seven capsule-deficient *K. pneumoniae* strains were calculated using three or more replicates for each condition; error bars represent standard error of the mean (SEM) measured by OD at 600 nm in liquid broth in the absence (grey) or presence (orange) of phage mtp5. At t = 0, the multiplicity of infection (MOI) was $10^2$.

**Figure S4.** Genetic structure of anti-$K^d$ phages mtp5 and mtp7 and the closest anti-Klebsiella phages available in public databases.

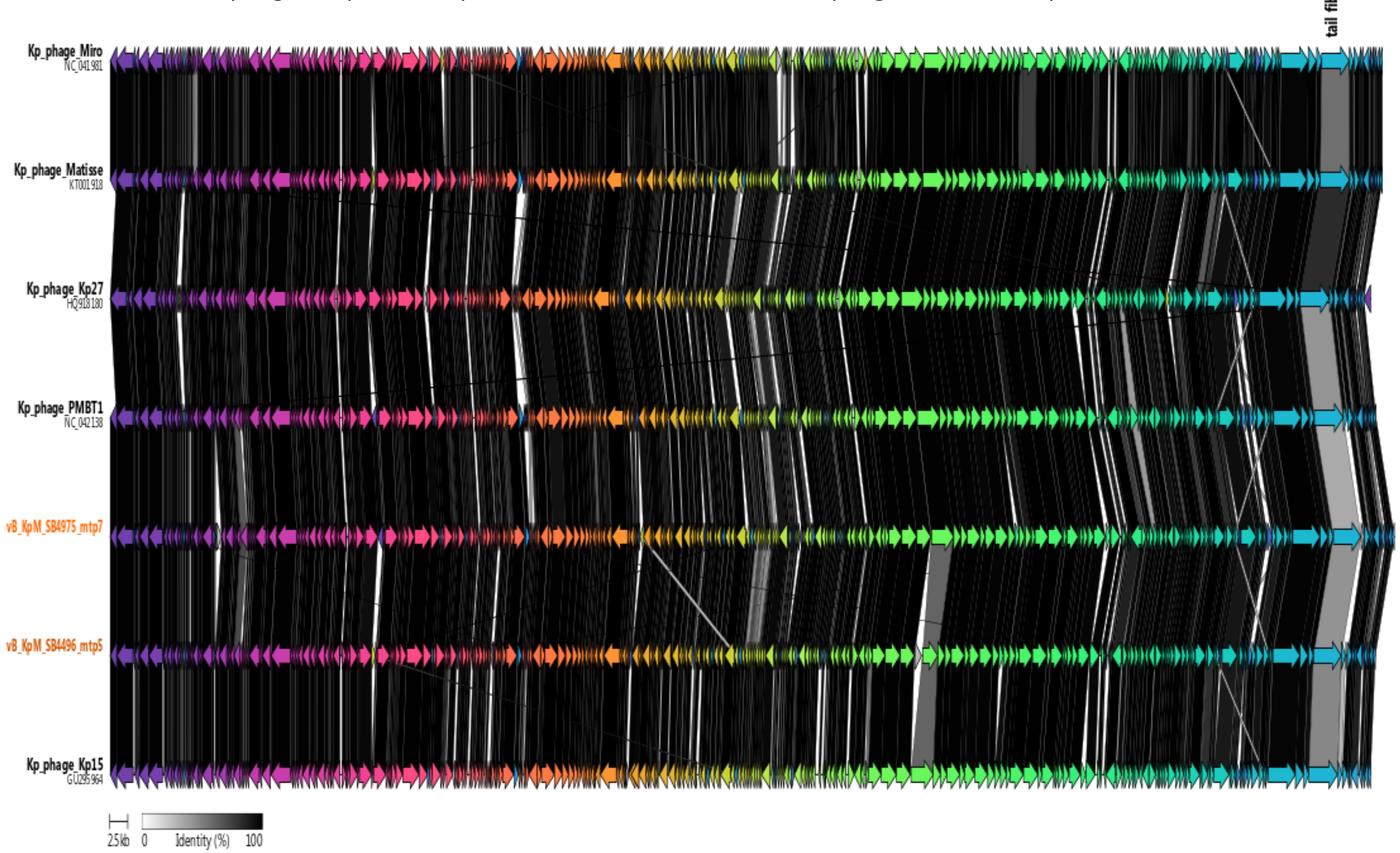


Phages Kp15, Kp27, Matisse, Miro and PMBT1 were identified based on BLASTN as the most similar to phage mtp5 and mtp7. Phages genomes were aligned using clinker. CDSs were arbitrarily colored. The white-to-dark grey blocks between CDSs represent their degree of AA identity (see color scale). ().

**Figure S5. Cocktails of phages mtp4, mtp6 and mtp7 with anti-capsule phages**

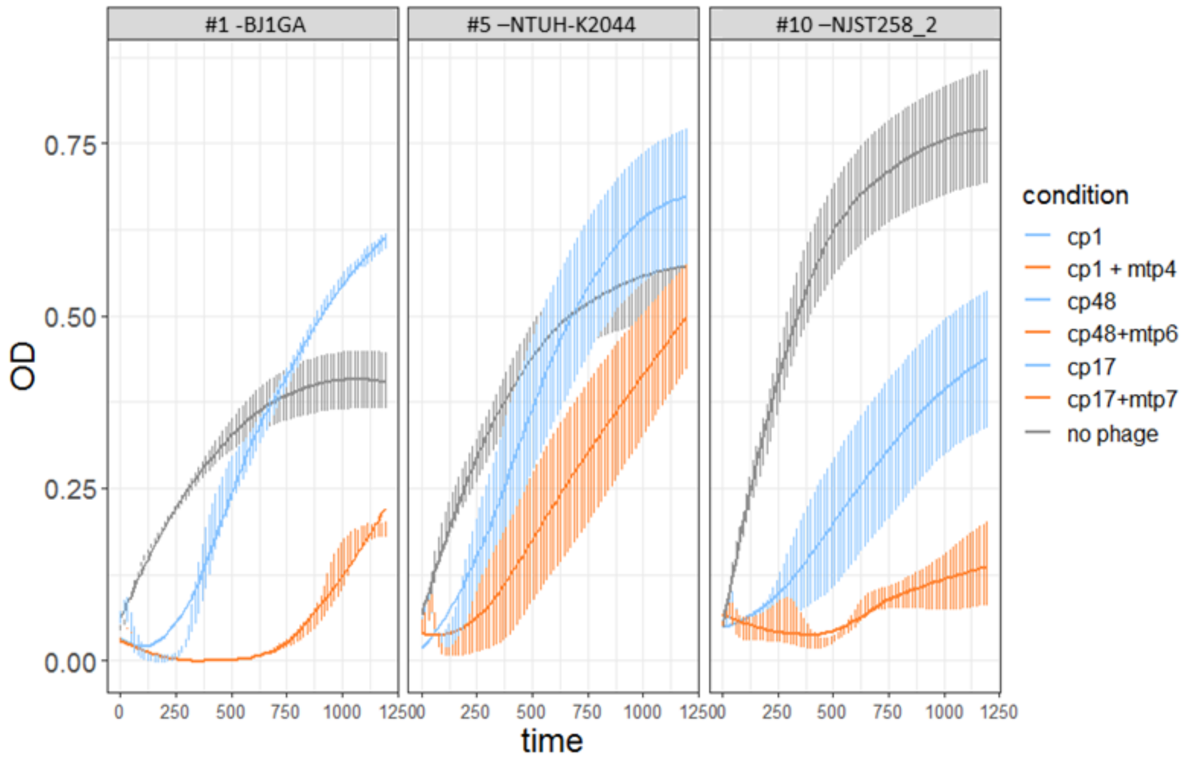


Growth curves for 3 different *K. pneumoniae* strains were obtained using three or more replicates for each condition; error bars represent standard error of the mean (SEM) measured via OD600 nm reading in liquid broth in the presence of three different phage cocktails. Cocktails containing anti-K phage and anti-$K^d$ phages are in orange, whereas the use of anti-K phage alone is depicted in blue (grey: control without phage). Left, anti-K phage cp1 and anti-$K^d$ phage mtp4 used against strain BJ1-GA; Center: anti-K phage cp17 and anti-$K^d$ phage mtp7 used against strain NJST258_2; Right: anti-K phage cp48 and anti-$K^d$ phage mtp6 tested against NTUH-K2044.

**Figure S6.** Subpopulations of bacteria that are non-susceptible to phages emerge more slowly when using anti-$K^d$ phages against non-capsulated strains

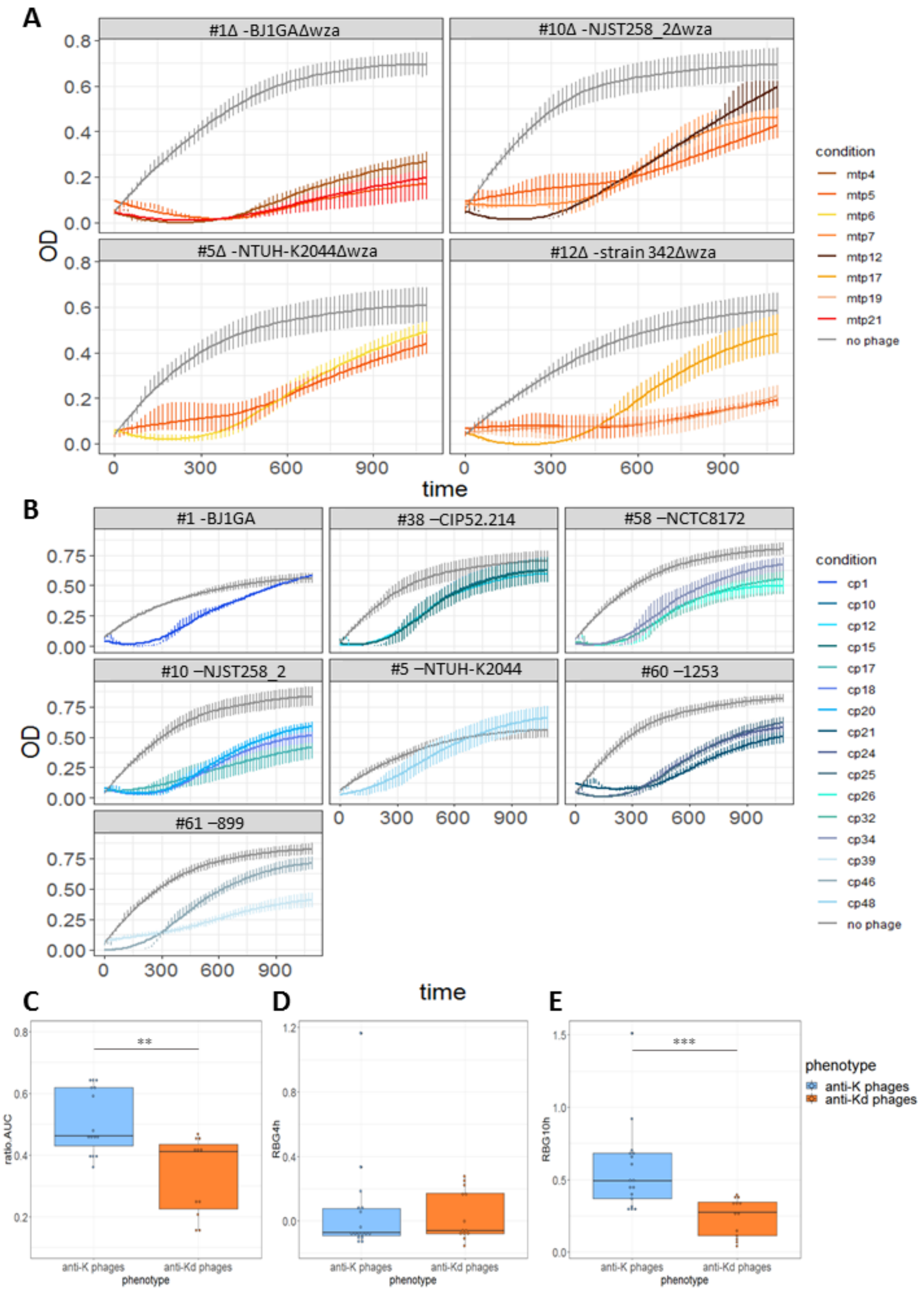

**A and B.** Growth curves obtained using three Kp strains with or without capsule. Three or more replicates were used for each condition; error bars represent standard error of the mean (SEM) based on OD600 nm reading in liquid broth in the absence (grey or presence of: **A.** anti-$K^d$ phages (orange tones) and **B.** anti-K phages (blue tones). At t = 0, multiplicity of infection (MOI) was $10^2$. In panels C, D and E, boxplots correspond to: **C.** Area under the curve (AUC); **D.** Relative bacterial growth (RBG) at 4h; and **E.** at 10h, of the time of regrowth on cultures targeted by anti-K phages vs anti-$K^d$ phages (**P-value ≤ 0.01;***P-value ≤ 0.001, Mann-Whitney test).

**Figure S7.** Resistance to anti-K phages leads to smaller (presumably less capsulated) colonies

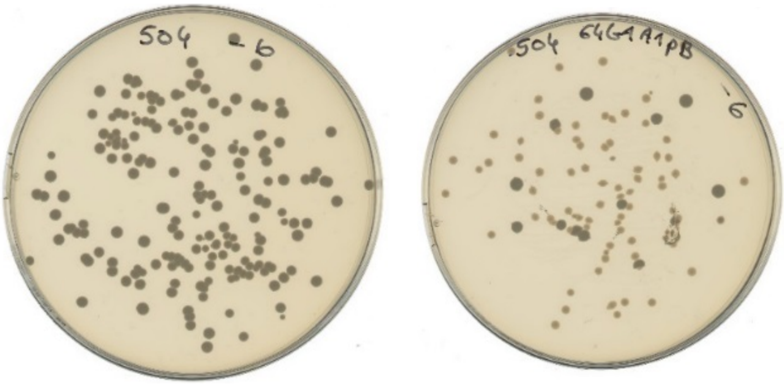

Example of appearance of non-capsulated colonies after anti-K phage use against capsulated strains. Here, culture of K64 strain NCTC8172, comparing the initial and the derived population, which was non-susceptible to phage cp34.

**Figure S8.** Mutations observed in populations exposed to phage cocktails that combine anti-K phages and anti-$K^d$ phages mtp4, mtp6 and mtp7

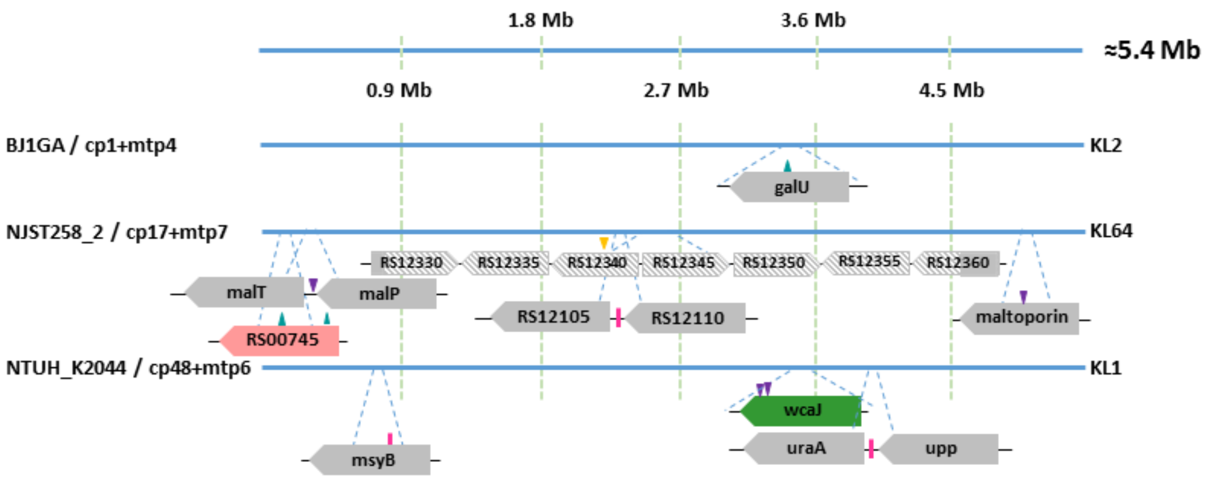


Genes colors: green: capsule locus; pink: LPS locus; and grey: others.

**Figure S9.** Phage mtp5 does not infect wild-type strain BJ1GA *in vitro* but infects it *in vivo*

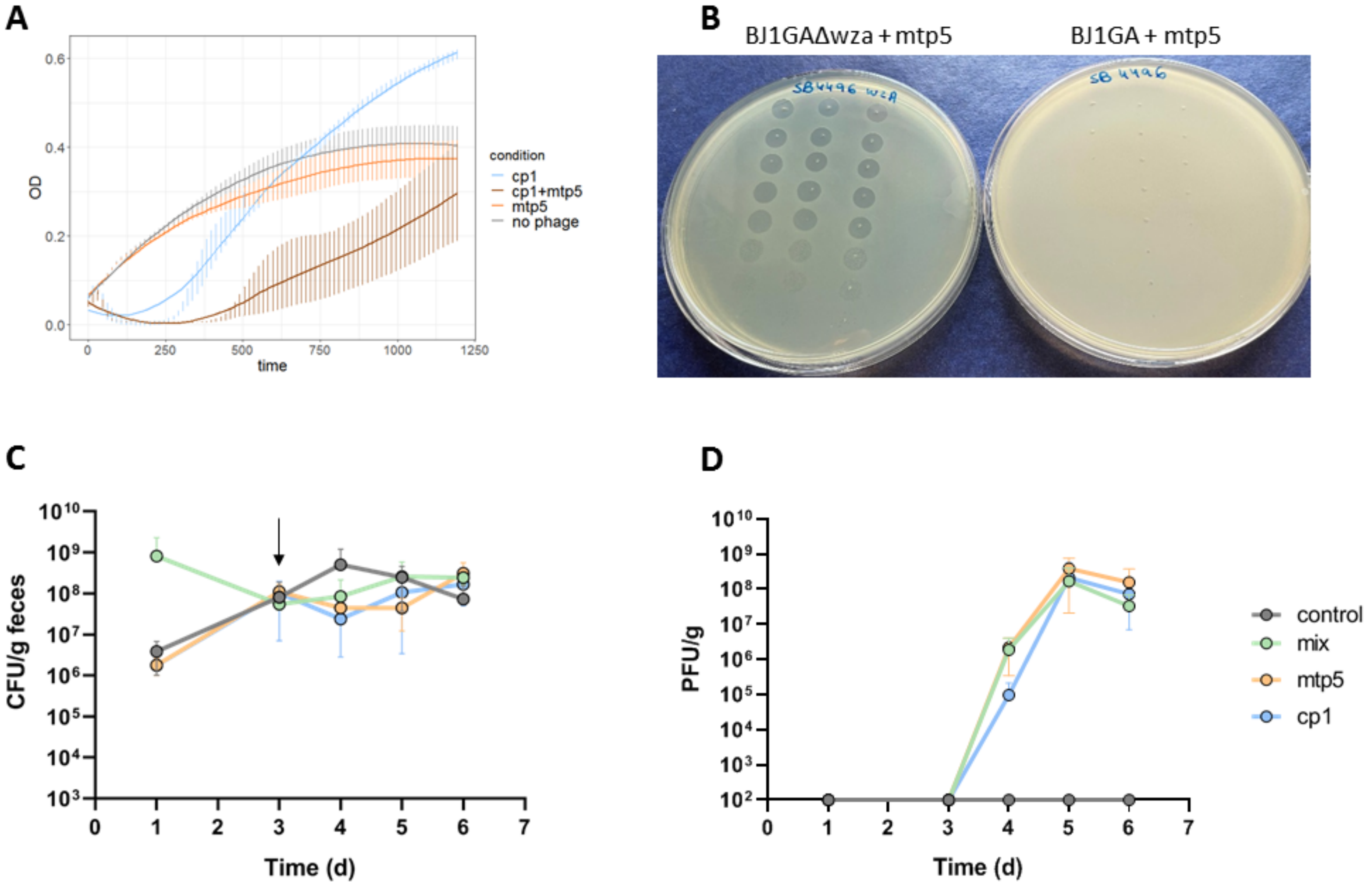

**A**: Growth curves measured by OD600 nm reading in liquid broth for strain BJ1GA in the presence or absence of phages cp1 or mtp5. **B**: Bacterial lawns spotted with both phage types, showing that mtp5 does not infect strain SB4496 in *in vitro* solid media conditions. **C and D**. *K. pneumoniae*-colonized OMM12 mice (n=13) received only at day 3 either PBS (pink, n=4), or the two phages cp1 and mtp5 together (mix; green, n=3; $6x10^7$ pfu per dose made of the same amount of each phage), or the individual phages cp1 (blue, n=3) and mtp5 (orange, n=3) by oral gavage. **C.** Levels of *K. pneumoniae* BJ1GA strain in the feces (the arrow indicates the day the phage was given to the mice). **D.** Phage titers from the fecal samples reported in panel C.

**Figure S10.** Isolation of different phenotypes from the feces of mice during phage treatment.

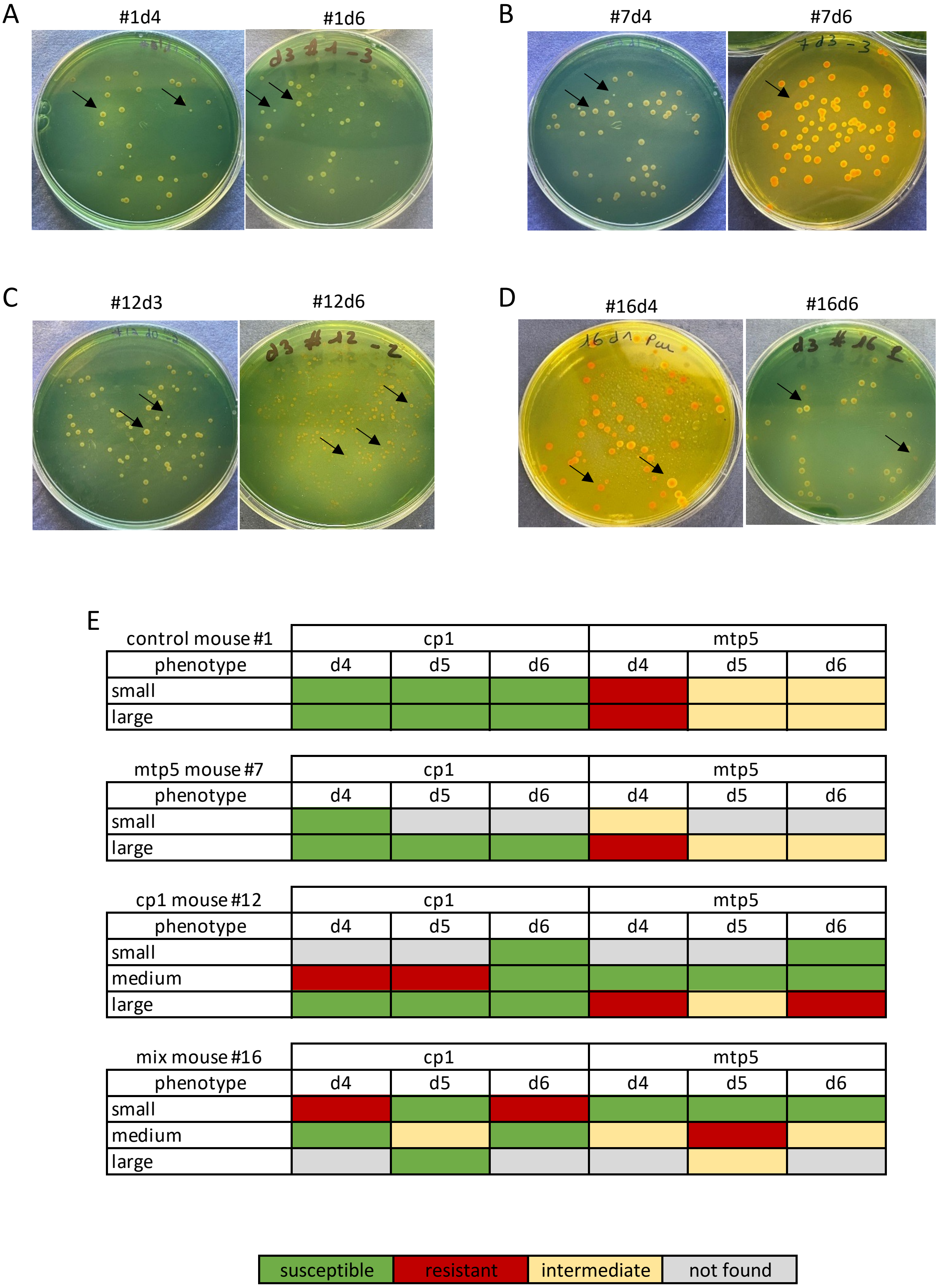

**A-D**. Examples of the colony phenotypes isolated from the feces of mice after K. pneumoniae colonization and phage treatment. **A.** Two K. pneumoniae phenotypes (small and large colonies) observed on SCAI medium agar from mouse #1 of the control group. **B.** Example of the two phenotypes isolated at day 4 (first day with phage) from feces of mouse #7 treated with phage mtp5 (the small phenotype was not observed the following days, at which only the large phenotype was observed . **C.** example of the three phenotypes isolated from feces of mouse #12 treated with phage cp1 (large, medium and small colony variants). **D.** Example of two of the three phenotypes isolated from feces of mouse #16 treated with cocktail of cp1+mtp5. **E.** All clones were tested for resistance against the two original phages, showing different susceptibility patterns.

**Figure S11.** Example of non-mucoid sectors

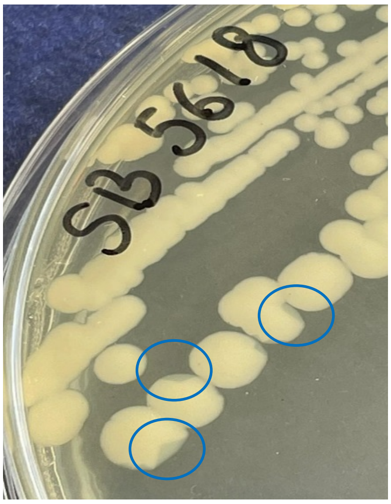


Capsulated (wild-type) *K. pneumoniae* strains were streaked on TSA (Tryptic Soy Agar) and incubated for 24 h at 37°C. The colonies were then searched for non-mucoid sectors; three examples are depicted on the plate. When none was detected, the plates were further kept at 25°C until non-mucoid sectors could be observed (Chiarelli et al. 2020).

**Figure S12.** Comparison of phages mtp6 and cp48 for their genomic structure and infection phenotype

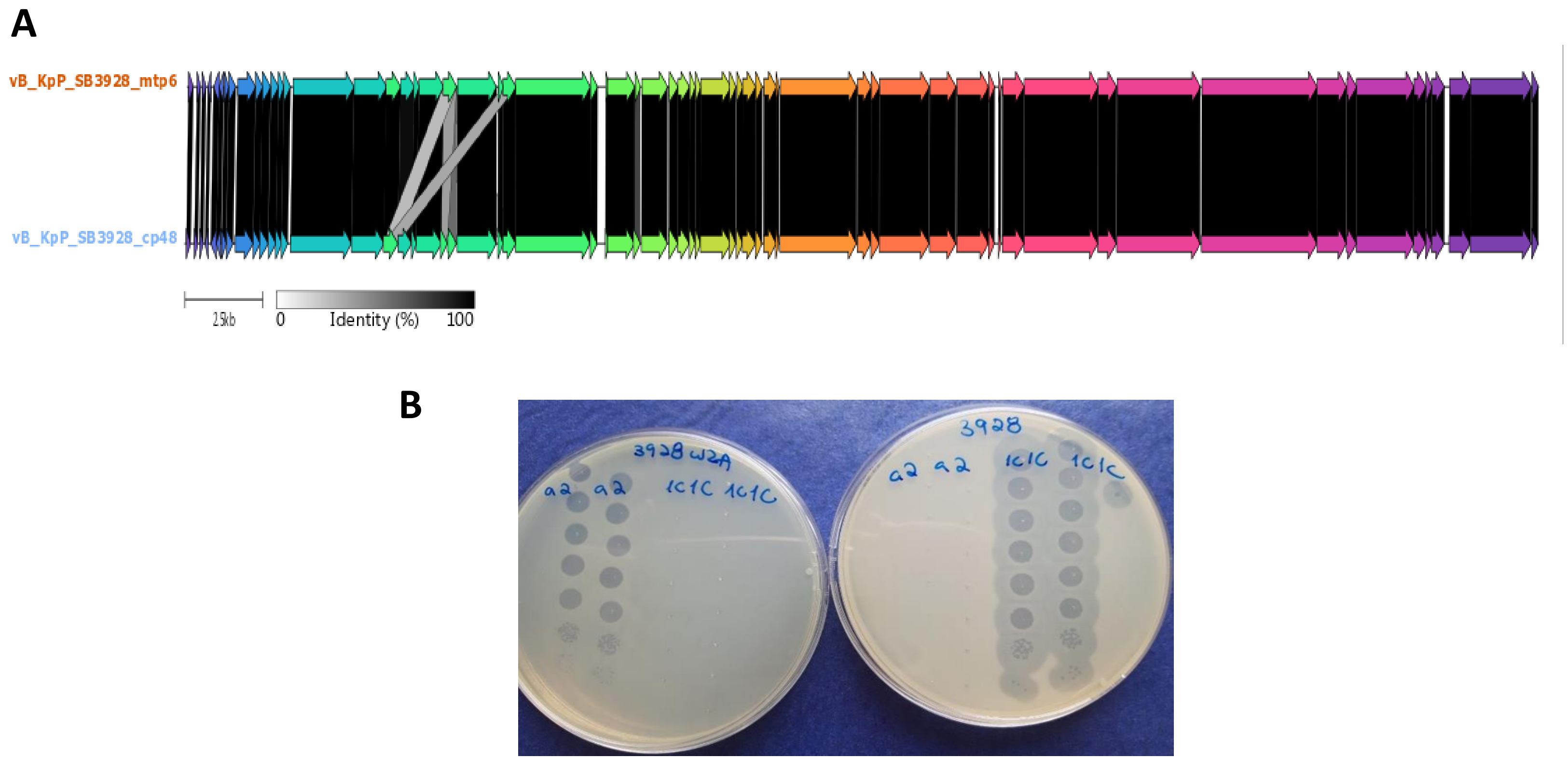


**A.** Comparison of the genomic structure of phages mtp6 and cp48. The only differences are observed in two hypothetical proteins and in one HNH endonuclease (see supplementary text2) B. Phenotypic differences in infection potential of phages mtp6 and cp48: whereas mtp6 infects the non-capsulated NTUK-2044 and cp48 the wt strain.

**Supplementary references**

Chiarelli, Adriana, Nicolas Cabanel, Isabelle Rosinski-Chupin, Pengdbamba Dieudonné Zongo, Thierry Naas, Rémy A. Bonnin, and Philippe Glaser. 2020. 'Diversity of Mucoid to Non-Mucoid Switch among Carbapenemase-Producing Klebsiella Pneumoniae'. *BMC Microbiology* 20 (1): 325. https://doi.org/10.1186/s12866-020-02007-y.

Deleo, Frank R., Liang Chen, Stephen F. Porcella, Craig A. Martens, Scott D. Kobayashi, Adeline R. Porter, Kalyan D. Chavda, et al. 2014. 'Molecular Dissection of the Evolution of Carbapenem-Resistant Multilocus Sequence Type 258 Klebsiella Pneumoniae'. *Proceedings of the National Academy of Sciences of the United States of America* 111 (13): 4988–93. https://doi.org/10.1073/pnas.1321364111.

Dong, Ning, Rong Zhang, Lizhang Liu, Ruichao Li, Dachuan Lin, Edward Wai-Chi Chan, and Sheng Chen. 2018. 'Genome Analysis of Clinical Multilocus Sequence Type 11 Klebsiella Pneumoniae from China'. *Microbial Genomics* 4 (2). https://doi.org/10.1099/mgen.0.000149.

Huynh, Bich-Tram, Virginie Passet, Andriniaina Rakotondrasoa, Thierno Diallo, Alexandra Kerleguer, Melanie Hennart, Agathe De Lauzanne, et al. 2020. 'Klebsiella Pneumoniae Carriage in Low-Income Countries: Antimicrobial Resistance, Genomic Diversity and Risk Factors'. *Gut Microbes*, May, 1–13. https://doi.org/10.1080/19490976.2020.1748257.

Kala, Smriti, Nichole Cumby, Paul D. Sadowski, Batool Zafar Hyder, Voula Kanelis, Alan R. Davidson, and Karen L. Maxwell. 2014. 'HNH Proteins Are a Widespread Component of Phage DNA Packaging Machines'. *Proceedings of the National Academy of Sciences of the United States of America* 111 (16): 6022–27. https://doi.org/10.1073/pnas.1320952111.

Lery, Letícia M. S., Lionel Frangeul, Anna Tomas, Virginie Passet, Ana S. Almeida, Suzanne Bialek-Davenet, Valérie Barbe, et al. 2014. 'Comparative Analysis of Klebsiella Pneumoniae Genomes Identifies a Phospholipase D Family Protein as a Novel Virulence Factor'. *BMC Biology* 12 (May): 41. https://doi.org/10.1186/1741-7007-12-41.

Merlet, A., C. Cazanave, H. Dutronc, B. de Barbeyrac, S. Brisse, and M. Dupon. 2012. 'Primary Liver Abscess Due to CC23-K1 Virulent Clone of Klebsiella Pneumoniae in France'. *Clin Microbiol Infect* 18 (9): E338-9. https://doi.org/10.1111/j.1469-0691.2012.03953.x.

Rodrigues, Carla, Siddhi Desai, Virginie Passet, Devarshi Gajjar, and Sylvain Brisse. 2022. 'Genomic Evolution of the Globally Disseminated Multidrug-Resistant Klebsiella Pneumoniae Clonal Group 147'. *Microbial Genomics* 8 (1). https://doi.org/10.1099/mgen.0.000737.

Sousa, Jorge A. M. de, Amandine Buffet, Matthieu Haudiquet, Eduardo P. C. Rocha, and Olaya Rendueles. 2020. 'Modular Prophage Interactions Driven by Capsule Serotype Select for Capsule Loss under Phage Predation'. *The ISME Journal* 14 (12): 2980–96. https://doi.org/10.1038/s41396-020-0726-z.

Turton, J. F., H. Englender, S. N. Gabriel, S. E. Turton, M. E. Kaufmann, and T. L. Pitt. 2007. 'Genetically Similar Isolates of Klebsiella Pneumoniae Serotype K1 Causing Liver Abscesses in Three Continents'. *J Med Microbiol* 56 (Pt 5): 593–97.

Wu, Keh-Ming, Ling-Hui Li, Jing-Jou Yan, Nina Tsao, Tsai-Lien Liao, Hui-Chi Tsai, Chang-Phone Fung, et al. 2009. 'Genome Sequencing and Comparative Analysis of Klebsiella Pneumoniae NTUH-K2044, a Strain Causing Liver Abscess and Meningitis'. *Journal of Bacteriology* 191 (14): 4492–4501. https://doi.org/10.1128/JB.00315-09.

Wyres, Kelly L., Jane Hawkey, Marit A. K. Hetland, Aasmund Fostervold, Ryan R. Wick, Louise M. Judd, Mohammad Hamidian, Benjamin P. Howden, Iren H. Löhr, and Kathryn E. Holt. 2019. 'Emergence and Rapid Global Dissemination of CTX-M-15-Associated Klebsiella Pneumoniae Strain ST307'. *The Journal of Antimicrobial Chemotherapy* 74 (3): 577–81. https://doi.org/10.1093/jac/dky492.
